# Systematic analysis uncovers SYK dependency in NF1^LoF^ melanoma cells

**DOI:** 10.1101/2022.03.06.483170

**Authors:** Cara Abecunas, Christopher Whitehead, Elizabeth Ziemke, Douglas G. Baumann, Christy Frankowski-McGregor, Judith Sebolt-Leopold, Mohammad Fallahi-Sichani

**Affiliations:** Department of Biomedical Engineering, University of Virginia, Charlottesville, VA 22908; Department of Biomedical Engineering, University of Michigan, Ann Arbor, MI 48109; Department of Radiology, University of Michigan Medical School, Ann Arbor, MI 48109; Department of Pharmacology, University of Michigan Medical School, Ann Arbor, MI 48109; Rogel Cancer Center, University of Michigan, Ann Arbor, MI 48109; UVA Comprehensive Cancer Center, University of Virginia, Charlottesville, VA 22908; Dewpoint Therapeutics, 451 D Street, Suite 104, Boston, MA 02210

**Keywords:** melanoma, NF1 loss of function, small-molecule kinase inhibitors, SYK kinase, mitochondrial electron transport chain, drug discovery, systems biology

## Abstract

The loss of function (LoF) of NF1 is the third most frequent mutation that drives hyperactivated RAS and tumor growth in >10% of melanomas. NF1^LoF^ melanoma cells, however, do not show consistent sensitivity to individual MEK, ERK, or PI3K/mTOR inhibitors. Here, we perform a targeted kinase inhibitor screen and identify a tool compound, named MTX-216, to be highly effective in blocking NF1^LoF^ melanoma cells. Single-cell analysis links drug-induced cytotoxicity to effective co-suppression of proliferation marker Ki-67 and the ribosomal S6 phosphorylation, an integrator of multiple RAS-mediated signaling pathways. Using a combination of kinome selectivity assay, transcriptomic analysis, and genetic experiments, we find the anti-tumor efficacy of MTX-216 to be dependent on its ability to inhibit not only PI3K (its nominal target) but also SYK, and suppression of a group of genes that regulate mitochondrial electron transport chain and whose expression is associated with poor survival in NF1^LoF^ melanoma patients. Furthermore, combinations of inhibitors targeting either MEK or PI3K/mTOR with an independent SYK kinase inhibitor or SYK knockdown show favorable effects. These studies provide a path to exploit SYK dependency to selectively block NF1^LoF^ melanoma cells.

**Statement of significance:** NF1^LoF^ melanomas represent a subtype with hyperactivated RAS signaling, for which currently no targeted therapies are clinically available. Our systems pharmacology studies identify SYK as a new vulnerability in NF1^LoF^ melanoma cells.

## Introduction

Neurofibromin 1 (NF1) is a tumor suppressor gene encoding a RAS GTPase-activating protein, whose loss of function (LoF) is the third most frequent mutation present in approximately 13% of melanomas (1–4). The loss of function of NF1 abrogates negative feedback on RAS, thereby causing hyperactivation of the downstream MAPK and PI3K/mTOR signaling pathways (5,6). NF1^LoF^ mutations, however, do not always predict sensitivity to MEK/ERK kinase inhibitors or PI3K/mTOR inhibitors and most NF1^LoF^ melanomas show only partial responses to these therapies (7–9).

Multiple strategies have been tested to enhance treatment efficacy in NF1^LoF^ melanomas. The most commonly explored approaches have been focused on testing combinations of small molecules (e.g., MEK/ERK inhibitors and PI3K/mTOR inhibitors) that target parallel pathways downstream of RAS signaling, in order to prevent the bypass mechanisms of drug resistance (10). Targeting RAS regulators that function downstream of receptor tyrosine kinases (RTKs) has been proposed as another strategy based on evidence that NF1^LoF^ cells require upstream RTK signals to drive aberrant cell growth (11). SHP2 inhibitors, for example, decouple the RAS/MAPK pathway from external growth signals and thereby reduce tumor growth in NF1^LoF^ melanoma cells (12). An alternative strategy would be to evaluate the effectiveness of multi-targeted inhibitors that block key signaling molecules upstream and downstream of RAS (13,14). A possible advantage of this approach may be a further increase in treatment efficacy that results from co-targeting the pathways on which NF1^LoF^ cell growth and survival are dependent as well as adaptive pathways that diminish the efficacy of individual MEK/ERK or PI3K/mTOR inhibitors.

In this paper, we perform a targeted compound screen to search for actionable kinases upstream, parallel to, or downstream of MAPK and PI3K/mTOR pathways, whose inhibition individually or in combination with one another efficiently block NF1^LoF^ melanoma cells. We identify MTX-216, a dual ATP-competitive inhibitor of PI3K isoforms and EGFR, to kill NF1^LoF^ melanoma cells in a dosedependent manner and suppress NF1^LoF^ melanoma tumor growth *in vivo.* Single-cell analysis reveals a substantial correlation between drug efficacy in NF1^LoF^ melanoma cells and effective co-suppression of Ki-67 (a proliferation marker) and the ribosomal S6 phosphorylation (an integrator of multiple RAS-mediated signaling pathways). These effects, however, are not explained merely by the nominal targets of MTX-216. Kinome selectivity analysis integrated with differential gene expression and pathway enrichment analysis identify SYK as an additional target of MTX-216 that is required for NF1^LoF^ melanoma cell growth. SYK is a non-receptor tyrosine kinase conventionally known as a mediator of adaptive immune receptor signaling (15). Recent studies, however, have implicated SYK localization to mitochondrial intermembrane space and its involvement in the regulation of mitochondrial respiratory chain and oxidative metabolism (16–20). Consistent with these studies, we find that MTX-216 treatment in NF1^LoF^ melanoma cells suppresses genes encoding mitochondrial electron transport chain. In agreement with these findings, treatment with IACS-010759, a clinical-grade small-molecule inhibitor of the mitochondrial electron transport chain (21), also induces cell death in these cells. Therefore, the selectivity profile of MTX-216, which includes SYK inhibition and the consequent suppression of mitochondrial electron transport chain, causes the unusually high activity of this compound in NF1^LoF^ melanoma cells.

## Materials and Methods

### Cell lines and reagents

Melanoma cell lines used in this study were obtained from the following sources: LOXIMVI (CVCL_1381) and COLO858 (CVCL_2005) from MGH Cancer Center, COLO792 (CVCL_1134) from ECACC, MeWo (CVCL_0445), Ramos Burkitt’s lymphoma (CVCL_0597) and A375 (CVCL_0132) from ATCC, SKMEL113 (CVCL_6074) from Memorial Sloan-Kettering Cancer Center, WM3918 (CVCL_C279) from Rockland Immunochemicals, human epidermal primary melanocytes (NHEM-Ad) from Lonza, YUHEF (CVCL_G326), YURED (CVCL_K020), YUHOIN (CVCL_J521) and YUTICA (CVCL_J071) from Yale University Dermatology Cell Culture Facility. All cell lines have been periodically subjected to re-confirmation by Short Tandem Repeat (STR) profiling by ATCC and mycoplasma testing by MycoAlert^TM^ PLUS mycoplasma detection Kit (Lonza). YUHEF, YURED, YUHOIN, and YUTICA cells were grown in Opti-MEM (Gibco, Cat# 31985088) supplemented with 5% fetal bovine serum (FBS). COLO858, LOXIMVI, COLO792, and SKMEL113 cells were grown in RPMI 1640 (Corning, Cat# 10-040-CV) supplemented with 1% Sodium Pyruvate (Invitrogen) and 5% FBS. A375 cells were grown in DMEM with 4.5 g/L glucose (Corning, Cat# 10-013-CV) supplemented with 5% FBS. MeWo cells were grown in EMEM (Corning, Cat# 10-009-CV) supplemented with 5% FBS. WM3918 cells were grown in 80% MCDB 153 (Sigma-Aldrich, Cat# M7403), 20% Leibovitz’s L-15 (Sigma-Aldrich, Cat# L1518), 5% FBS, and 1.68 mM CaCl_2_. Ramos cells were grown in RPMI 1640 supplemented with 1% sodium pyruvate and 10% FBS. NHEM-Ad cells were grown in MGM-4 Melanocyte Growth Medium-4 BulletKit (Lonza, Cat# CC-3249). We added penicillin (50 U/mL) and streptomycin (50 lg/mL) to all growth media.

Small molecule inhibitors trametinib, ulixertinib, AZD8055, palbociclib, gefitinib, pictilisib, entospletinib, R406 and IACS-010759 were all purchased from Selleck Chemicals. MTX-216 and MTX-211 were synthesized by Albany Molecular. MTX-211 (HY-107364) is also available from MedChemExpress. Compounds used for cell-based studies were dissolved in the appropriate vehicle (DMSO) at a stock concentration of 5 or 10 mM.

The following primary monoclonal antibodies (mAb) and polyclonal antibodies (pAb) with specified animal sources, catalogue numbers and dilution ratios, were used in immunofluorescence staining and Western blot assays: Ki-67 (AB_2797703) (mouse mAb, Cell Signaling Technology, Cat# 9449, 1:800), p-S6^S235/S236^ (AB_10695457) (rabbit mAb, Cell Signaling Technology, Cat# 4851, 1:400), p-p44/42 MAPK (ERK1/2) (T202/Y204) (AB_2315112) (rabbit mAb, Cell Signaling Technology, Cat# 4370, 1:800), p-AKT^S473^ (AB_2315049) (rabbit mAb, Cell Signaling Technology, Cat# 4060,1:400), p-4EBP1^T37/46^ (AB_560835) (rabbit mAb, Cell Signaling Technology, Cat# 2855, 1:200), SYK (AB_628308) (mouse mAb, Santa Cruz Biotechnology, Cat# 1240, 1:1000), p-SYK^Y525/526^ (AB_2197222) rabbit mAb, Cell Signaling Technology, Cat# 2710, 1:1000), p-BLNK^Y96^ (AB_2064915) (rabbit pAb, Cell Signaling Technology, Cat# 3601, 1:1000), BLNK (AB_626752) (mouse mAb, Santa Cruz Biotechnology, Cat# 8003, 1:1000), and β-tubulin (AB_2210545) (rabbit pAb, Cell Signaling Technology, Cat# 2146, 1:1000). The following primary mAb and pAb antibodies with specified animal sources, catalogue numbers and dilution ratios, were used in Western blot assays in xenograft pharmacodynamic (PD) studies: SYK (AB_2687924) (rabbit mAb, Cell Signaling Technology, Cat# 13198, 1:1000), EGFR (AB_2230881) (rabbit mAb, Cell Signaling Technology, Cat# 2646, 1:1000), AKT (AB_329827) (rabbit pAb, Cell Signaling Technology, Cat# 9272, 1:1000), β-actin (AB_10950489) (rabbit mAb, Cell Signaling Technology, Cat# 8457, 1:1000), p-SYK^Y525/526^ (rabbit mAb, Cell Signaling Technology, Cat# 2710, 1:1000), p-EGFR^Y1068^ (AB_2096270) (rabbit mAb, Cell Signaling Technology, Cat# 3777, 1:1000), and p-AKT^S473^ (rabbit mAb, Cell Signaling Technology, Cat# 4060, 1:1000). The following secondary antibodies with specified sources and catalogue numbers were used at 1:2000 dilution for immunofluorescence assays: anti-mouse Alexa Fluor 488 (AB_141607) (Thermo Fisher, Cat# A21202) and anti-rabbit Alexa Fluor 647 (AB_2536183) (Thermo Fisher, Cat# A31573). The following secondary antibodies with specified sources and catalogue numbers were used at 1:15000 dilution for Western blot assays: anti-rabbit IRDye 800CW (AB_621848) (LI-COR Biosciences, Cat# 926-32213) and anti-mouse IRDye 680RD (AB_10953628) (LI-COR Biosciences, Cat# 926-68072). The following secondary antibody with specified source and catalogue number was used at 1:10000 dilution for Western blot PD assays: horseradish peroxidase-conjugated anti-rabbit antibody (AB_2313567) (rabbit IgG, Jackson ImmunoResearch Laboratories Inc., Cat# 111-035-003).

### High-throughput drug treatment and measurements of drug sensitivity

Growth rate inhibition assays with kinase inhibitors were performed in 96-well clear bottom black plates (Corning, Cat# 3904). Viable cells were counted using a TC20 automated cell counter (SCR_008426) (Bio-Rad Laboratories) and seeded in 200 μL of full growth media at a density of 1,700-2,500 cells per well. Cells were treated the next day with small molecule compounds at reported concentrations or with vehicle control (DMSO) using a D300e Digital Dispenser. At designated timepoints (t = 24, 72, 120 and 168 h), surviving cells were fixed, stained with Hoechst 33342 (Thermo Fisher, Cat# H3570), imaged and counted following image analysis as described in the immunofluorescence staining and image analysis section.

The net growth rate for cells treated with individual compounds (at different doses), or their combination, or vehicle, was calculated from time-dependent changes in live cell count according to the following equation:

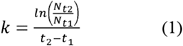

where *N_t1_* and *N_t2_* represent the number of cells measured at timepoints *t* = *t_1_* and *t* = *t_2_,* respectively, and *k* describes the net growth rate of cells during the time period between *t_1_* and *t_2_*. The net growth rates were then averaged across multiple, consecutive time intervals to determine the mean net growth rate for each condition. To compare drug sensitivity among different cell lines, we normalized the mean net growth rate measured for drug-treated cells (*k*_drug_) to that measured for vehicle (DMSO)-treated cells (*k*_DMSO_) to calculate the drug-induced normalized growth rate (a.k.a. DIP rate (22)):

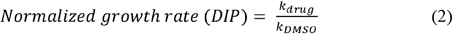

Area over the dose-response curve (AOC) (23) was calculated across all tested inhibitor concentrations (*c_i_*) as follows:

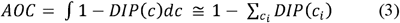

### Immunofluorescence staining and image analysis

Cells in Corning 96-well plates were fixed with 4% PFA for 20 min at room temperature, washed twice with PBS, and then permeabilized in 100% ice-cold methanol for 10 min at −20° C. Cells were rewashed with PBS, blocked using Odyssey Blocking Buffer (LI-COR) for 1 h at room temperature, and stained with primary antibodies at their appropriate dilutions in Odyssey Blocking Buffer overnight (~16 h) at 4° C. The next day, cells were washed three times with PBS supplemented with 0.1% Tween-20 (Sigma-Aldrich) (PBS-T), and then stained with secondary antibodies for 1 h at room temperature followed by washing once with PBS-T, once in PBS, and incubation with Hoechst 33342 for 20 min at room temperature. Cells were washed twice with PBS and imaged with a 10x objective using the ImageXpress Micro Confocal High-Content Imaging System (Molecular Devices) or the Operetta CLS High-Content Imaging System (Perkin Elmer). A total of 9 sites were imaged per well. Background subtraction was performed with ImageJ v.2.1.0 software (SCR_003070). Image segmentation and signal intensity quantification was performed with CellProfiler v.3.1.9 (SCR_007358) (24). Single-cell data were analyzed using MATLAB 2019b (SCR_001622). By generating histograms of single-cell data across a variety of conditions for each protein (X), including positive and negative controls, we identified an appropriate binary gate, based on which the percentage of X^High^ versus X^Low^ cells in each condition was quantified.

### NucView apoptosis assay

COLO792 and MeWo cells were seeded in 96-well clear bottom black plates in full phenol red free growth media (100 μL/well). WM3918 were seeded in their usual full growth media. Cells were treated the next day with indicated concentrations of MTX-216 (in duplicate) using D300e Digital Despenser. After 48 or 72 h of treatment, a reagent cocktail comprised of Hoechst 33342 and NucView 488 Caspase-3 reporter (Biotium, Cat# 30029) was added to each well for final concentrations of 1 μg/mL and 1 μM, respectively. The plates were then incubated in a tissue culture incubator (37°C, 5% CO2) for 1.5 h. Pre-warmed PFA was then added to each well for a final concentration of 1%. Plates were then sealed using Microseal aluminum foil (Bio-Rad), incubated for 30 min at room temperature and imaged with a 10x objective. Image analysis and quantification was performed as described in the immunofluorescence staining and image analysis section.

### Annexin V and propidium iodide (PI) double-staining apoptotic assay

WM3918 cells were harvested 48 h after treatment with inhibitor or DMSO. FITC Annexin V/Dead Cell Apoptosis Kit (Invitrogen, Cat# V13242) was used to stain cells for Annexin V and propidium iodide (PI). As per the manufacturer’s protocol, samples were washed with cold PBS and suspended in 1x annexin binding buffer. Cell suspensions were mixed with 5 μL FITC Annexin V and 1 μL PI (100 μg/mL) for 15 min at room temperature. The mixture was further diluted with 1x annexin binding buffer. Samples were run in BD LSRFortessa™ Cell Analyzer (SCR_018655) (BD Biosciences). Flow data was cleaned up by removing debris and analyzed in FCS Express 7 Research (SCR_016431) (DeNovo Software).

### Biochemical kinase inhibition assays

To assess their ability to inhibit each of 403 human kinases, MTX-216 and MTX-211 were outsourced to SelectScreen Biochemical Kinase Profiling service at ThermoFisher. Each compound was initially screened at 10 μM using either Z’-LYTE or Adapta kinase activity assay at an ATP concentration of K_m,app_ for each target. The Z’-LYTE assay monitors phosphorylation of peptide substrates, and Adapta quantifies ADP formation and thus can be used with lipid substrates. For selected targets that were inhibited significantly (>90%) by MTX-216 or MTX-211 at 10 μM (e.g., SYK), the kinase activity assay was repeated across 10 concentrations. Dose-response plots and the corresponding IC_50_’s were generated using the function doseResponse (developed by Ritchie Smith) available on the MATLAB Central File Exchange (https://www.mathworks.com/matlabcentral/fileexchange/33604-doseresponse).

### Gene knockdown by siRNA

siRNA knockdown experiments with MeWo and YUHEF cells were performed in antibiotic-free growth media either in cells seeded in 10-cm plates (for the purpose of Western blotting) or in Corning 96-well clear bottom black plates (at a density of 2,500 and 1,000 cells/well, respectively) for the purpose of cell counting and inhibitor sensitivity analysis. In both cases, 24 h after cell seeding, cells were transfected with Dharmacon’s ON-TARGETplus Human SMARTpool siRNA for SYK (L-003176-00-0005), nontargeting control (D-001810-10-05), or the four SYK-specific siRNAs that make up the SYK SMARTpool (J-003176-10, J-003176-11, J-003176-12, J-003176-13) at 25 nM each. DharmaFECT 2 (GE Dharmacon, Cat# T-2002-01) and DharmaFECT 3 (GE Dharmacon, Cat# T-2003-01) were used as transfection reagents for MeWo and YUHEF cells, respectively. After 72 h of culture in the presence of siRNAs, cells in 10-cm plates were transferred to ice, washed with ice-cold PBS, and lysed for the purpose of Western blotting. After 72 h of knockdown, cells in 96-well plates were either fixed with 4% PFA or treated with kinase inhibitors (trametinib, MTX-211, pictilisib) or DMSO for an additional 72 h before fixation. All fixed cells were then stained with Hoechst 33342 and counted using quantitative immunofluorescence microscopy. Time-dependent changes in cell count were used to compute normalized growth rates as described above.

### *In vivo* xenograft assays

Animal handling, care and treatment was carried out in accordance with procedures approved by the Institutional Animal Care and Use Committee (IACUC) at the University of Michigan. For the COLO792 xenograft studies, female 5-6 weeks old NCR nude mice (IMSR_TAC:ncrnu) (Taconic, CrTac:NCr-*Foxn1^nu^*) were implanted subcutaneously with 2.5×10^6^ cells in 1:1 Matrigel:serum-free media. When the average tumor volume was approximately 125 mm^3^ (on day 39 post-implantation), mice were randomized into treatment groups and treatments were initiated. MTX-216 was prepared fresh daily as a clear solution in 1:2 propylene glycol:1% Tween80/Na_3_PO_4_, and administered intraperitoneally (IP) based upon individual animal body weight (0.2 mL/20 grams) on days 1 (i.e., day 39 post-implantation), 2, 4, 7, 10, 13 and 16. Tumor volumes were measured on days 1, 3, 6, 8, 12, 15, 18, and 20 post-treatment. For the WM3918 xenograft studies, female 5-6 weeks old NCR nude mice (Taconic) were implanted subcutaneously with 1.5×10^6^ cells in 1:1 Matrigel:serum-free media. When the average tumor volume was approximately 115 mm^3^ (on day 6 post-implantation), mice were randomized into treatment groups and treatments were initiated. MTX-216 was prepared fresh daily as a fine, light brown homogenous suspension including 5% DMSO and 95% PBS with a pH of 7. MTX-216 was administered once daily intraperitoneally (IP) based upon individual animal body weight (0.2 mL/20 grams), beginning on day 1 (i.e., day 6 post-implantation). Tumor volumes were measured on days 1, 3, 5, 7, 9 and 11. Tumor volumes were calculated by measuring two perpendicular diameters with calipers and using the formula: tumor volume (mm^3^) = (length × width^2^)/2. Animal body weights were measured two to three times a week. At the completion of the experiment or when vehicle-treated tumor volume exceeded 1,000 mm^3^, the mice were euthanized.

To measure the pharmacodynamic effect of MTX-216 *in vivo,* female 5-6 weeks old NCR nude mice (Taconic) were implanted subcutaneously with 3×10^6^ WM3918 cells in 1:1 Matrigel:serum-free media. When tumors reached approximately 150 mm^3^, mice were randomized into treatment groups. Mice received a single treatment of vehicle control (5% DMSO/95% PBS), MTX-216 at 50 mg/kg, or MTX-216 100 mg/kg IP. At 30 minutes and 60 minutes post treatment, mice were euthanized, and the tumors were harvested and snap frozen in liquid nitrogen.

### Western blots

Cultured melanoma cells treated with indicated siRNAs or inhibitors were transferred to ice, washed with ice-cold PBS, and lysed with ice-cold RIPA buffer (Thermo Fisher, Cat# 89901) consisting of 25 mM Tris-HCl (pH 7.6), 150 mM NaCl, 1% NP-40, 1% sodium deoxycholate, and 0.1% SDS supplemented with 1 mM Na3VO4, 15 mM NaF, cOmplete ULTRA EDTA-free protease inhibitors (Roche, Cat# 5892953001) and PhosSTOP (Roche, Cat# 4906837001). Protein samples of cleared lysates were measured using BCA assay (Thermo Scientific, Cat# PI23235). Lysates were adjusted to equal protein concentrations for each treatment condition, 4× NuPage LDS sample buffer (Invitrogen, Cat# NP0007) supplemented with 1 μL of 1 M DTT was added, and samples were heated for 10 min at 70°C. Samples were then loaded on NuPAGE 3-8% Tris-Acetate gels (Invitrogen). Western blots were performed using iBlot 2 Gel Transfer Stacks PVDF system (Invitrogen). Membranes were blocked with Intercept (TBS) Blocking Buffer (LI-COR, Cat# 927-60001) for 1 h and then incubated with primary antibodies (1:1000 dilution) in 0.5x Intercept blocking buffer solution (diluted with TBS, supplemented with 0.1% Tween-20) overnight at 4° C. Secondary antibodies were used at 1:15000 dilution in 0.5x Intercept blocking buffer solution supplemented with 0.01% SDS. Membranes were scanned on an Odyssey CLx scanner (LI-COR) with 700 and 800 nm channels set to automatic intensity at 169 μm resolution. Blot images were processed, on Image Studio Lite v.5.2.5 software (LI-COR).

Ramos cells were treated with indicated inhibitors for 6 h prior to stimulating the cells with 25 μg/mL IgM (Goat F(ab’)2 Anti-Human IgM, Southern Biotech, Cat# 2022-01) or BBS control for 3 min. Samples were lysed, processed, and run in Western blots as described above.

To measure the pharmacodynamic effect of MTX-216 on p-SYK^Y525/526^, p-EGFR^Y1068^, p-AKT^S473^ *in vivo,* excised xenograft tumors were rinsed in cold 1X PBS before adding complete RIPA lysis buffer. Tumors were homogenized using a douncer and rocked for 1 hour at 4°C. Tumors were then centrifuged at 4°C for 20 minutes at 13.2 rpm. Quantitation of protein lysates was performed using DC Protein Assay (BioRad) according to the manufacturer’s instructions. Lysates were adjusted to equal protein concentrations using water, 1X LDS Sample Buffer (Life Technologies), and 1 M DTT. Samples were heated for 10 min at 70°C and loaded into NuPage 4-12% Bis-Tris gel. Proteins were transferred to a 0.45 μm PVDF membrane (Immobilon-P) for 50 minutes at 20V in a semi-dry transfer apparatus containing Transfer Buffer. Membranes were blocked for 1 hour at room temperature in 5% BSA TBS-T or 5% NFDM TBS-T. Blocked membranes were incubated overnight at 4°C with primary antibodies (p-SYK^Y525/526^, p-EGFR^Y1068^, p-AKT^S473^, SYK, EGFR, AKT, β-actin) in 5% BSA TBS-T or 5% NFDM TBS-T, as recommended by the manufacturer. Membranes were incubated for 1 hour at room temperature in horse radish peroxidase-conjugated anti-rabbit antibody in 1% BSA TBS-T or 1% NFDM TBS-T. The membranes were incubated in ECL Prime Western Blotting Detection Reagents (Cytiva) for 30 seconds before exposure to autoradiography film.

### MitoTracker staining, imaging and quantitation

Cells were seeded at 100 μL/well in 96-well PerkinElmer ViewPlates (PerkinElmer, Cat# 6005225) or 96-well Cellvis glass coverslip bottom plates (Cellvis, Cat# P96-1.5P). Cells were treated the next day with either SYK siRNAs (or non-targeting control), 0.1-1 μM MTX-216, 0.1-1 μM MTX-211, or DMSO in triplicate. At the designated timepoints (72 h in case of siRNA knockdown and 48 h in case of inhibitor treatments), MitoTracker Deep Red (Invitrogen, Cat# M22426) and MitoTracker Red CMXRos (Invitrogen, Cat# M7512) were added to the wells for a final concentration of 100 nM for each dye. The plates were then incubated in a tissue culture incubator (37°C, 5% CO2) for 45 min. The media including dye solution were aspirated, wells were washed with pre-warmed media (including inhibitor or DMSO) and placed in incubator for 1 h to wash out nonspecific binding of dye. Cells were then washed with PBS followed by 4% PFA fixation for 15 min at room temperature. Plates were washed three times (5 min each) with PBS followed by Hoechst 33342 staining for 20 min. Plates were washed twice with PBS and sealed using Microseal aluminum foil (Bio-Rad). PerkinElmer plates were imaged with 20x water-immersion objective and Cellvis glass plates were imaged with 20x air objective using the Operetta CLS High-Content Imaging System (Perkin Elmer). A total of 37 sites were imaged per well. Image analysis was performed as described in the immunostaining and image analysios section.

### RNA extraction, library construction, and RNA sequencing analysis

MeWo, WM3918, and SKMEL113 cells were seeded in 10-cm plates in three replicates, and treated the next day with either DMSO, MTX-216 (at 1 μM), MTX-211 (at 1 μM), pictilisib (at 1 μM), trametinib (at 0.1 μM), or the combination of MTX-216 (at 1 μM) and trametinib (at 0.1 μM), for 24 h. At the time of harvest, cells were washed once with PBS and then lysed with Rneasy Buffer RLT (Qiagen) in the dish. Lysate samples were then vortexed and frozen. To extract RNA, lysate samples were thawed at room temperature and then processed using QIAshredder (Qiagen, Cat# 79654) and Rneasy Mini Kit (Qiagen, Cat# 74104). RNA was Dnase treated (Qiagen, Cat# 79254) and then eluted. RNA quality was assessed by Bioanalyzer (Agilent). All samples had RINs of 9.0 or higher. Prior to poly-A enrichment, 500 ng of each RNA sample was supplemented with 1 μL (1:100 dilution) of ERCC spike-in control (Life Technologies, Cat# 4456740). Poly-A RNA enrichment was performed using NEBNext Poly(A) kit (New England BioLabs). Fragments were end-modified with adaptors using NEBNext Multiplex Oligos for Illumina Set 2 (New England BioLabs), following library amplification (PCR, 11 cycles). Resulting libraries were prepared using NEBNext Ultra II Directional RNA Kit (New England BioLabs). Samples were run on Illumina NovaSeq 6000 (SCR_01638) (S4) using paired-end read with 150 cycles. In total, 1.4 billion reads were generated, including ~33 million reads per sample. Reads were mapped to the human genome (hg38) using HISAT2 (SCR_015530) (25), transcripts were assembled and quantified using StringTie (SCR_016323) (26,27), and differential gene expression analysis was performed by DESeq2 (SCR_015687) using the web-based Galaxy platform (SCR_006281) (28,29).

To identify genes regulated uniquely by MTX-216 versus MTX-211 in WM3918 and MeWo cells, we first selected those genes with base mean ≥ 1, that changed in abundance by more than twofold relative to a DMSO-treated control (|log_2_(fold change)ļ ≥ 1; FDR < 0.05) in each cell line. Among these genes, we then identified the subset that was similarly up- or down-regulated by each compound in both cell lines. Genes enriched specifically by MTX-216 treatment (but not in MTX-211 treatment) in both cell lines were used in Enrichr (SCR_001575) (https://maayanlab.cloud/Enrichr/) (30) and compared statistically with gene expression signatures extracted from the Gene Expression Omnibus (GEO) (SCR_005012) database for kinase perturbation (31).

To systematically perform differential gene expression analysis across three cell lines treated with different compounds (MTX-216, MTX-211, pictilisib, trametinib, MTX-216 plus trametinib, or DMSO), we used principal component analysis (PCA). For PCA, we first organized the gene expression dataset into a matrix with 42 rows (representing 3 cell lines, 6 distinct treatment conditions, and 3 independent replicates) and 16,940 columns (representing genes). The list of these genes was generated after filtering out transcripts with unidentified gene names and functions in StringTie and removing those genes for which all 42 conditions had TPM values < 1. Then log_2_(TPM+1) values were calculated for each gene and z-scored across all 42 conditions. PCA was then performed using the MATLAB built-in function pca.

Because PC3 and PC4 separated the efficacious inhibitors independent of cell lines, we used PC3 and PC4 loadings to identify genes associated with distinct inhibitory effects on cells treated with different compounds. Gene loadings that were differentially enriched along either PC3 or PC4 were selected based on a ratio cutoff of *cotangent* (π/8) ≈ 2.4. Among these genes, we identified the subsets along each PC that had loading values of greater than 1.5×median for positive loadings, or smaller than 1.5×median for negative loadings. These genes were then used in Enrichr to identify significant Gene Ontology (GO) Biological Processes (SCR_002811).

### Hierarchical clustering

Following PCA, unsupervised hierarchical clustering of expression levels of genes with negative PC3 or PC4 loadings was carried out using MATLAB 2020b with the Euclidean distance metric and the Average linkage algorithm for computing the distance between clusters.

### The Cancer Genome Atlas (TCGA) bioinformatics analysis

The clinical history and the corresponding gene expression (STAR count) data from patients enrolled in the Skin Cutaneous Melanoma (SKCM) project of the TCGA (1) were accessed using the cBio Cancer Genomics Portal (32) (http://cbioportal.org/) and the Genomic Data Commons Data Portal (SCR_014514) (https://portal.gdc.cancer.gov/). TCGA-SKCM samples were classified based on their NF1 mutation status. NF1^LoF^ samples were those bearing putative driver “truncating” mutations. NF1^Mutant^ samples were those bearing putative driver “splice” or “missense” mutations. NF1^Other^ samples were those bearing no NF1 mutations or putative “passenger” mutations. Sequencing reads were accessed through *GDCquery* function in the *TCGAbiolinks* package (SCR_017683) in R (SCR_001905) (33). A total of 469 sequenced melanoma tumors were analyzed: NF1^LoF^ (39 samples), NF1^Mutant^ (9 samples), and NF1^Other^ (421 samples). RNA-seq reads were then normalized to adjust for GC-content effects on read counts and distributional differences between samples using *TCGAanalyze_Normalization* function within the *TCGAbiolinks* package. Genes were then z-scored across all tumors. We then calculated mean z-score values across the genes found within Respiratory Electron Transport Chain GO term (GO:0022904) (http://geneontology.org/). Statistical analysis of ETC expression of NF1^LoF^ patients versus NF1^Mutant^ or NF1^Other^ was analyzed in R using the built-in function *t.test*. Tumors for each NF1 cohort were then stratified into two subgroups, ETC^High^ and ETC^Low^, based on the on the top and bottom quartiles. We then used the tumors with ETC^High^ and ETC^Low^ signatures in NF1^LoF^ and NF1^Other^ cohorts to compare the overall patient survival between each subgroup based on two-sided *P* values computed by log-rank (Mantel-Cox) test. Stage of diagnosis comparing ETC^High^ and ETC^Low^ within the NF1^LoF^ subgroup was analyzed using the Neoplasm Disease Stage American Joint Committee on Cancer Code database, followed by Chi-squared test and Benjamini-Hochberg FDR correction. SYK mutation analysis across TCGA-SKCM was also analyzed with Chi-squared test followed by Benjamini-Hochberg FDR correction. All statistical results were calculated and provided by cBio Cancer Genomics Portal.

### Gene set enrichment analysis

Gene set enrichment analysis (GSEA) (SCR_003199) (34,35) was performed using GSEA v.4.1.0 or v.4.2.3 software. For the pre-rank gene analysis, the GSEAPreranked module was used. To analyze the drug effects across the entire Hallmark database (36), genes were pre-ranked based on their log_2_-fold change of expression (by MTX-216 or trametinib relative to DMSO) and *P* values generated from DESeq2 for three replicates in each of the three cell lines tested (MeWo, WM3918 and SKMEL113) using the following equation:

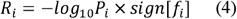

where *R_i_* represents the rank for gene *i*, *P_i_* represents the *P* value associated with change of expression for gene *i*, and *f_i_* represents the log2 fold change associated with gene *i*. GSEA was performed with 1,000 permutations across Hallmark gene sets to determine downregulated and upregulated biological pathways.

To assess the effects of individual drugs compared to DMSO with respect to MAPK and PI3K/mTOR signaling pathways, we ran pre-rank gene analysis using the log2 fold change generated from DESeq2. GSEAPreranked was performed with 1,000 permutations across GO ERK1 and ERK2 Cascade (GO:0070371) and Hallmark PI3K AKT MTOR Signaling gene signatures.

To compare SYK activity in NF1^LoF^ or NF1 Mutant versus NF1-Other patient samples, normalized TCGA-SKCM sequenced samples were run against eachother using the default GSEA parameters with 1,000 permutations. Genes downregulated due to SYK drug inhibition identified in Figure 3 from the GEO kinase perturbation database were used in the enrichment anlaysis (i.e., SYK_druginhibition_282_GSE43510, SYK_druginhibition_286_GSE43510_downregulated) (30,31).

### Statistics and reproducibility

No statistical method was used to predetermine sample size. Sample sizes were chosen based on similar studies in the relevant literature. The experiments were not randomized. The investigators were not blinded to allocation during experiments and outcome assessment. All data with error bars were presented as mean values ± s.d. or mean values ± s.e.m. as indicated in figure legends using indicated numbers of independent replicates. The significance of pairwise correlations was evaluated based on *P* values associated with the corresponding two-sided Pearson’s correlation analysis (*r* = correlation coefficient). Comparisons between groups were performed using the Student’s *t* test or ANOVA as described in figure legends. Statistical analyses were performed using MATLAB 2020b software.

### Data availability

All data generated in this study are included in this manuscript or its supplementary information files and the raw sequencing datasets have been deposited into Gene Expression Omnibus (GEO): GSE213588. Data analysis was performed in MATLAB and R using built-in functions and parameters as described in Methods.

### Materials Availability

All cell lines and reagents used in this study are available via commercial or non-commercial resources described.

## Results

### MTX-216 blocks NF1^LoF^ melanoma cell growth *in vitro* and *in vivo*

To identify kinases whose inhibition might elicit efficacious anti-tumor responses in NF1^LoF^ melanoma cells, we assembled a panel of small-molecule inhibitors targeting different kinases that act downstream or upstream of RAS signaling, including: MEK1/2 inhibitor (trametinib), ERK1/2 inhibitor (ulixertinib), pan-PI3K inhibitor (pictilisib), mTORC1/2 inhibitor (AZD8055), CDK4/6 inhibitor (palbociclib), and EGFR inhibitor (gefitinib). We also tested the impact of inhibiting multiple kinases, either by using the combination of two distinct compounds (e.g., ulixertinib plus palbociclib) or by using multi-kinase inhibitors such as the dual PI3K/EGFR inhibitor, MTX-216, and its structural analogue, MTX-211 (Fig. 1A, Supplementary Fig. S1A). We profiled the effects of these compounds on cell growth and survival in a panel of nine NF1^LoF^ melanoma cell lines. Among the tested cell lines, six expressed a wild-type (WT) BRAF (BRAF^WT^/NF1^LoF^), two possessed BRAF^V600E^/NF1^LoF^ double mutations, and one carried NRAS^Q61R^/NF1^LoF^ double mutations (Supplementary Fig. S1B). Each inhibitor was tested over a range of four concentrations, including a maximum dose (C_max_) of approximately 10 times the reported biochemical IC_50_ for its nominal target. To compare inhibitor sensitivities, we calculated the normalized growth rates (22) by dividing the 7 day-averaged net growth rate for cells treated with each inhibitor (at each dose) to net growth rate for vehicle (DMSO)-treated cells (Fig. 1B, Supplementary Fig. S1C). Normalized growth rates < 0 indicated a net cell loss (i.e., cytotoxicity), a value of 0 represented no change in viable cell number (i.e., cytostasis), and a value of 1 represented no drug effect. All the examined inhibitor conditions led to variable responses across NF1^LoF^ cell lines. MTX-216 was, however, consistently cytotoxic, leading to steep dose-response curves. The maximal effect (E_max_) of MTX-216 was significantly higher than that of other tested compounds, including MAPK inhibitors trametinib and ulixertinib, which exhibited potent but less efficacious responses in NF1^LoF^ cell lines (Fig. 1B, C). Among the tested inhibitor combinations, MTX-216 plus trametinib or ulixertinib was most effective, leading to efficacious and potent responses (Supplementary Fig. S1D, E), suggesting that MTX-216 and MAPK inhibitors may target orthogonal vulnerabilities in NF1^LoF^ melanoma cells.

**Figure 1.**
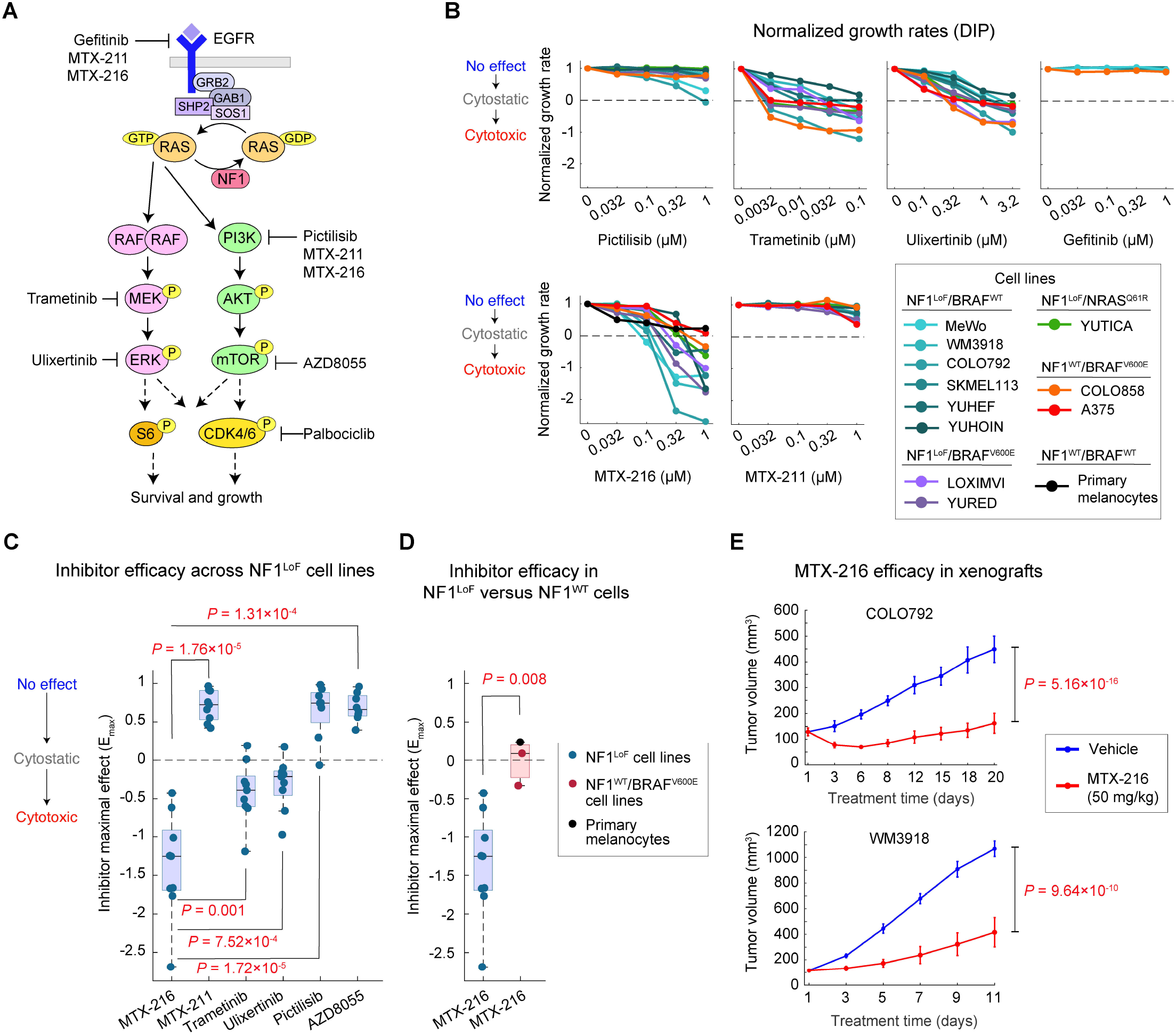
MTX-216 blocks tumor growth in NF1^LoF^ melanomas *in vitro* and *in vivo.* **(A)** A simplified schematic illustration of the function of NF1 in the context of RAS-mediated activation of ERK and PI3K/mTOR signaling. The nominal targets for kinase inhibitors tested in this study are highlighted. **(B)** Dose-dependent changes in normalized growth rates (a.k.a. DIP rates) across cell lines with different NF1/BRAF/NRAS mutation status or primary epidermal melanocytes. The average net growth rates for each condition were calculated from measurements of live cell count (across two replicates) at four timepoints (including 1, 3, 5, and 7 days). **(C, D)** Inhibitor maximal effect (E_max_), defined as the normalized growth rate evaluated at the highest tested concentration, of different inhibitors across NF1^LoF^ melanoma cell lines **(C)** and between NF1^LoF^ cell lines and NF1^WT^ cells, including two BRAF^V600E^ melanoma cell lines and primary melanocytes **(D)**. Statistical significance was determined by two-sided, paired-sample *t* test **(C)** or two-sided, two-sample *t* test **(D)**. **(E)** Mice bearing xenografts of COLO792 cells (top) and WM3918 cells (bottom) were treated with MTX-216 or vehicle for a period of 20 and 11 days, respectively, to determine the effect on tumor growth. Tumor volume data represent mean values ± s.e.m across 4 mice per group (in case of COLO792) or 5 mice per group (in case of WM3918). Statistical significance was determined by two-way ANOVA.

To determine whether the efficacy of MTX-216 was specific to NF1^LoF^ cells, we evaluated its effect on non-transformed primary melanocytes as well as on two NF1^WT^/BRAF^V600E^ melanoma cell lines (A375 and COLO858) that are sensitive to trametinib and ulixertinib. When tested at its highest concentration (1 μM), MTX-216 caused no cytotoxicity in primary melanocytes (Fig. 1B, D). It also exhibited a substantially less efficacious response in A375 and COLO858 cells in comparison with any of the nine NF1^LoF^ cell lines tested (Fig. 1B, D).

We then asked whether sensitivity to MTX-216 was recapitulated *in vivo.* Athymic nude mice were subcutaneously implanted with either a slowly or rapidly growing NF1^LoF^ cell line, COLO792 or WM3918, respectively. Animals were then treated with MTX-216 (at a dose of 50 mg/kg) or vehicle by intraperitoneal injection (IP). Inhibitor administration in mice was performed every 1 to 3 days for a period of up to 20 days, during which body weight was also monitored. MTX-216 suppressed tumor growth in both cell line-derived xenograft models (Fig. 1E), while animals experienced no major loss in body weight (Supplementary Fig. S1F). Together, these data identify MTX-216 as an efficacious agent that selectively blocks NF1^LoF^ melanoma cells *in vitro* and shows encouraging signs of *in vivo* efficacy. We thus set out to identify the mechanisms that might determine the anti-tumor efficacy of MTX-216 in NF1^LoF^ melanomas.

### MTX-216 co-suppresses Ki-67 and p-S6 and induces apoptosis in NF1^LoF^ cells

To identify factors influencing the efficacy of MTX-216, we assayed apoptosis in NF1^LoF^ cells using two distinct methods. First, we exposed MeWo, WM3918 and COLO792 cells to increasing concentrations of MTX-216 for 48 or 72 h. We used the NucView® dye, a fluorogenic substrate of pro-apoptotic effector caspases 3 and 7, and measured substrate cleavage by fluorescence microscopy (37). Single-cell analysis revealed a dose-dependent increase in the frequency of apoptotic cells (with high caspase activity) induced by MTX-216 in all three NF1^LoF^ cell lines (Fig. 2A, B). We further confirmed the induction of apoptosis using a double-staining assay with Annexin-V and propidium iodide (PI) with flow cytometry. 48 h treatment of WM3918 cells with MTX-216 led to a ~3-fold increase in the proportion of early-stage apoptotic (Annexin-V^High^/PI^Low^) cells and ~4-fold increase in the proportion of late-stage apoptotic and necrotic (Annexin-V^High^/PI^High^) cells in comparison with DMSO-treated samples (Supplementary Fig. S2A).

**Figure 2.**
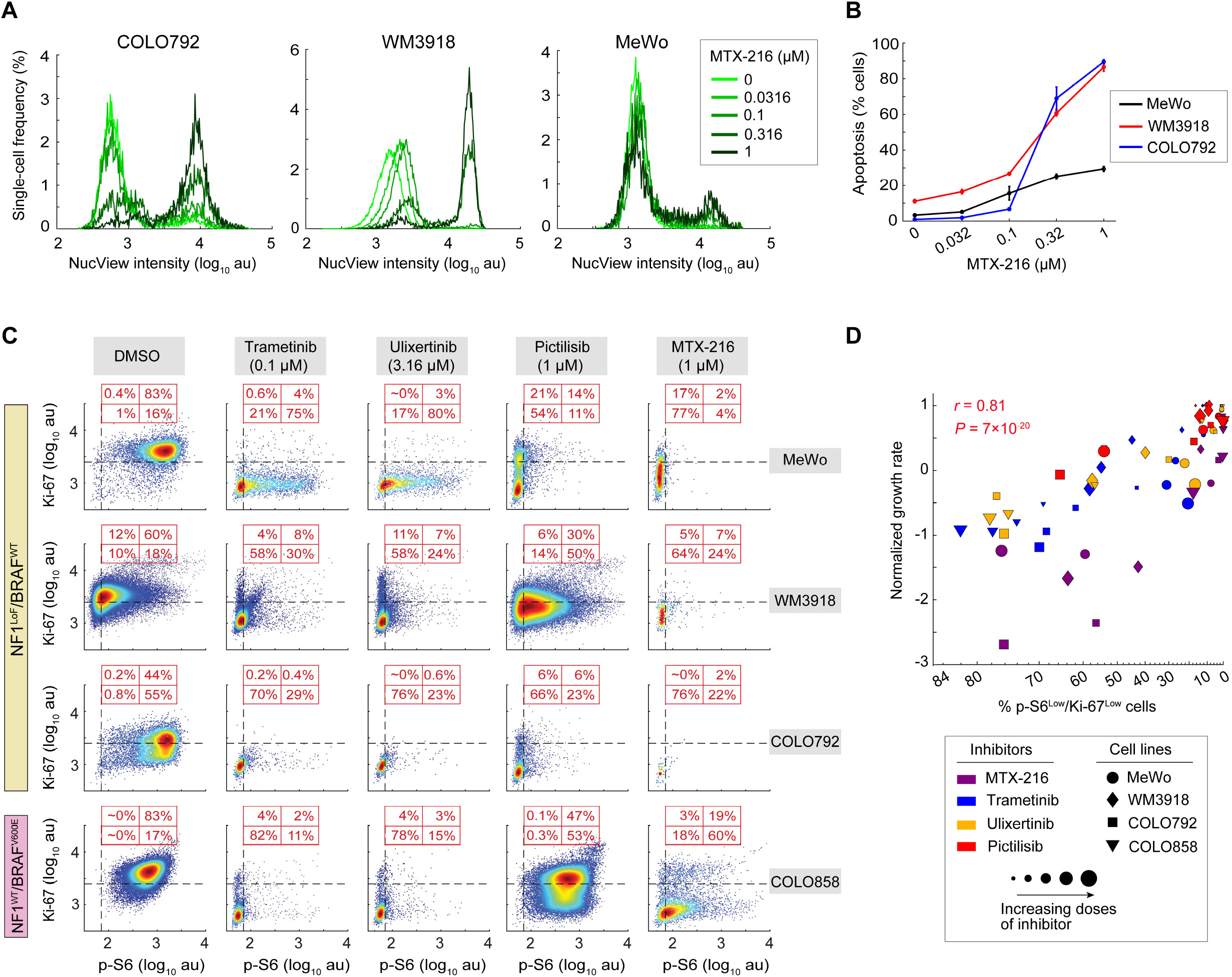
MTX-216 co-suppresses Ki-67 and p-S6 and induces apoptosis in NF1^LoF^ cells. **(A, B)** MTX-216 dose-dependent changes in apoptotic response of COLO792 cells (after 48 h) and WM3918 and MeWo cells (after 72 h), measured using a NucView Caspase-3/7 reporter assay. Single-cell NucView intensities across different inhibitor doses **(A)** and the estimation of the percentage of apoptotic cells based on NucView intensity gating **(B)** are shown for each cell line. Data are presented as mean values ± s.d. calculated across n = 2 replicates. **(C)** Co-variate single-cell analysis of p-S6 versus Ki-67 across three NF1^LoF^/BRAF^WT^ melanoma cell lines (MeWo, WM3918 and COLO792) and one NF1^WT^/BRAF^V600E^ cell line (COLO858) treated for 72 h with indicated concentrations of MTX-216, trametinib, ulixertinib and pictilisib. The horizontal and vertical dash lines were used to gate Ki-67^High^ versus Ki-67^Low^ cells and p-S6^High^ versus p-S6^Low^ cells, respectively. Percentages of cells in each quadrant are shown in red. **(D)** Two-sided Pearson’s correlation analysis between the percentage of p-S6^Low/^Ki-67^Low^ cells and the inhibitor-induced normalized growth rate (measured after 72 h of treatment) across three NF1^LoF^/BRAF^WT^ melanoma cell lines (MeWo, WM3918 and COLO792) and one NF1^WT^/BRAF^V600E^ cell line (COLO858) treated with MTX-216, trametinib, ulixertinib and pictilisib (at the same concentrations as shown in Fig. 1B).

To evaluate the effect of MTX-216 on signaling and proliferation, we quantified dose-dependent changes in the phosphorylation state of S6 ribosomal protein (p-S6^S235/S236^) and expression of Ki-67 protein by immunofluorescence microscopy. p-S6 serves as a surrogate for S6 kinase activity, integrating multiple pro-survival signals downstream of RAS signaling, including MEK/ERK and PI3K/mTOR pathways (38). Ki-67 is a marker of proliferation, which is expressed at lower levels in quiescent cells than in actively proliferating cells (39). We analyzed the single-cell expression of p-S6 and Ki-67 and their covariance in three NF1^LoF^ cell lines (MeWo, WM3918 and COLO792) following 72 h treatment with either MTX-216, pictilisib, trametinib or ulixertinib. All inhibitors reduced the expression of both markers (Fig. 2C and Supplementary Fig. S2B). MTX-216, however, was the only compound that consistently drove the largest proportion of NF1^LoF^ cells toward a doubly inhibited p-S6^Low^/Ki-67^Low^ state in all three NF1^LoF^ cell lines. In contrast, NF1^WT^/BRAF^V600E^ COLO858 cells (which show higher sensitivity to MEK/ERK inhibitors than to PI3K/mTOR inhibitors) exhibited more effective co-suppression of p-S6 and Ki-67 in response to trametinib and ulixertinib in comparison with MTX-216 (Fig. 2C).

Among all treatment conditions, the percentage of p-S6^Low^/Ki-67^Low^ cells significantly correlated with treatment-induced changes in growth rate (Fig. 2D). The high efficacy of MTX-216 in blocking NF1^LoF^ melanoma cells was, therefore, associated with its strong ability to co-suppress Ki-67 and p-S6 in these cells. To test whether such high efficacy could be explained by the compound’s direct effect on MAPK or AKT/mTOR pathways, we measured changes in p-ERK^T202/Y204^, p-AKT^S473^, and p-4EBP1^T37/46^ in MeWo cells following 6 h treatment with different concentrations of MTX-216 using quantitative immunofluorescence microscopy. For comparison, we also treated the cells with other inhibitors of the PI3K/mTOR and MAPK pathways, including pictilisib, AZD8055, MTX-211, and trametinib (Supplementary Fig. S2C, D). Trametinib was the only compound that reduced ERK phosphorylation in a dose-dependent manner, while MTX-211, MTX-216, pictilisib and AZD8055 had no significant effect on p-ERK levels. On the other hand, pictilisib, AZD8055, MTX-216, and MTX-211, but not trametinib, decreased p-AKT levels in a dose-dependent manner. The mTOR inhibitor AZD8055 was the only compound that potently reduced the levels of p-4EBP1, a marker of mTOR activity, while MTX-216, MTX-211 and pictilisib did not decrease p-4EBP1 levels at concentrations below 1 μM, and trametinib had no effect on p-4EBP1 levels. Overall, these data confirm the on-target effect of MTX-216 on the PI3K/AKT pathway and suggest that its high efficacy (in comparison with its structural analogue MTX-211 and other compounds tested in Figure 1) may be driven by mechanisms other than its direct nominal targets.

### MTX-216 suppresses the activity of SYK kinase

To identify what is driving the high levels of efficacy following MTX-216 treatment, kinome selectivity testing was carried out against 403 human kinases using two commercial FRET-based kinase assays, including Z’-LYTE which monitors phosphorylation of peptide substrates, and Adapta that quantifies ADP formation and can be used with lipid substrates. We also tested MTX-211 (the structural analogue of MTX-216) to quantify the differences in target space between the two compounds that might be responsible for their dramatically distinct effects on NF1^LoF^ cells. In the first step, our goal was to filter out kinases that were not inhibited by either compound at a high concentration (i.e., 10 μM). At this concentration, both compounds inhibited their nominal targets, including different isoforms of the catalytic subunits of PI3K (p110α, p110β, p110δ and p110γ) and EGFR, by approximately 90-100% (Fig. 3A). The assay also identified additional kinase targets inhibited by both compounds at 10 μM, including mTOR, PI4KB and DNA-PK. Repeating the assay for these targets across a ~10^4^-fold range of concentration showed that they were all inhibited by both compounds to similar degrees (i.e., similar IC_50_’s ranging from ~0.3 nM to 1 μM) (Supplementary Fig. S3A, Fig. 3B). We hypothesized that the superior efficacy of MTX-216 might be due to inhibition of a kinase that is targeted effectively by MTX-216 but not by MTX-211. The most significant candidate was SYK, a non-receptor tyrosine kinase, which was inhibited 94% and 52% by MTX-216 and MTX-211 at 10 μM, respectively (Fig. 3A). Comparing the kinase activity of SYK in the presence of various concentrations (from 0.5 nM to 10 μM) of each compound, we found that MTX-216 inhibited SYK with an IC_50_ of 281 nM, which was ~21-fold more potent than MTX-211 (with an IC_50_ of ~6 μM) (Fig. 3C).

**Figure 3.**
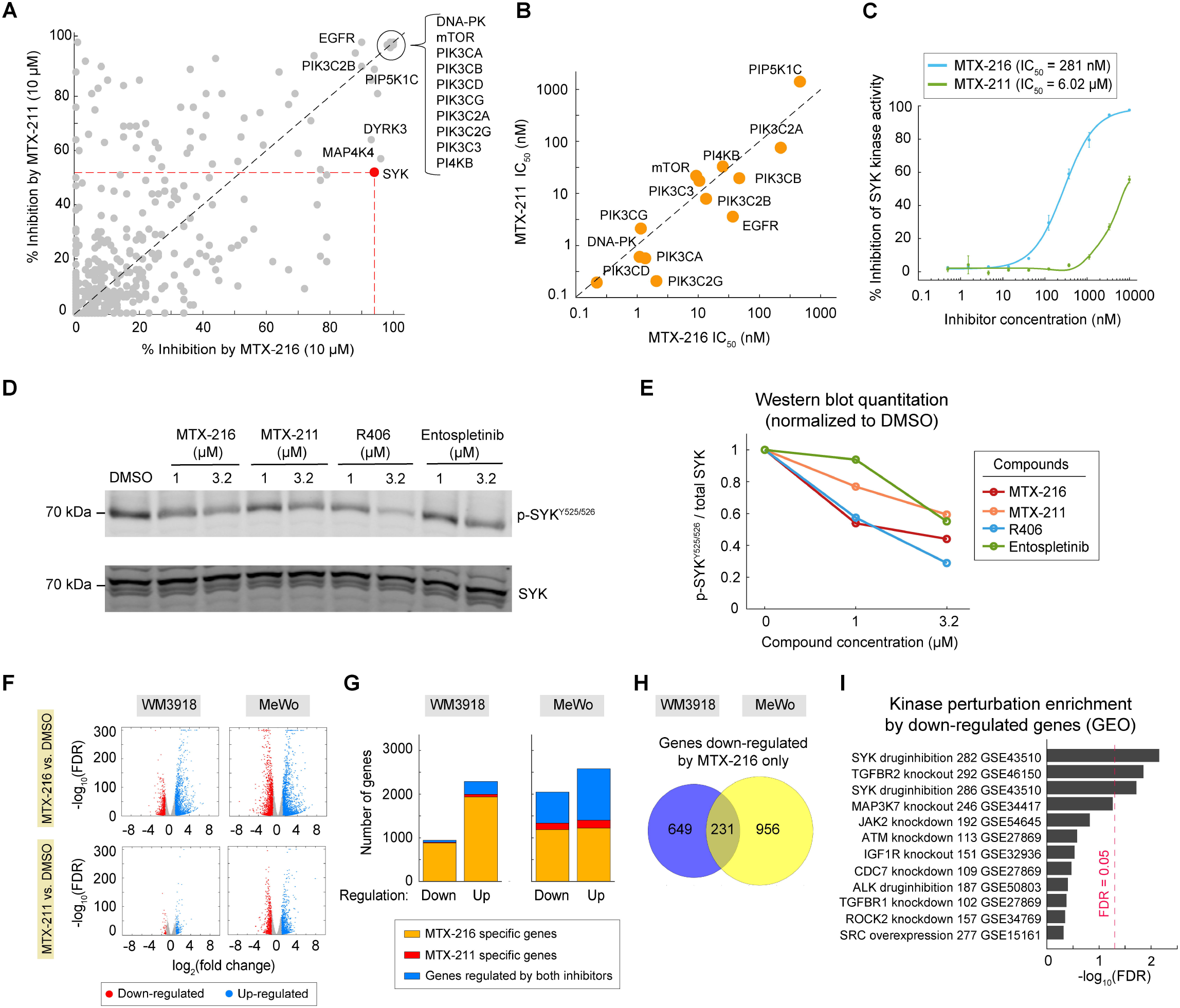
SYK kinase activity is more effectively inhibited by MTX-216 than by MTX-211. **(A)** High-throughput kinase inhibition assays (using Z’-LYTE and Adapta platforms) reporting % inhibition of 403 kinases by MTX-216 and MTX-211 (both at 10 μM) in the presence of ATP at a concentration of K_m_,_app_ for each target. Dashed line represents *y* = *x.* **(B)** Half-maximal inhibitory concentration (IC_50_) values for the nominal targets of MTX-216 and MTX-211 (PI3K and EGFR) as well as other kinases that were inhibited significantly (>90%) by both compounds (at 10 μM). IC_50_ values were derived from kinase activity assays performed across 10 concentrations of each compound. Dashed line represents *y* = *x*. **(C)** Dose-dependent inhibition of SYK kinase activity by MTX-216 and MTX-211. SYK kinase inhibition was profiled using Z’-LYTE assay in the presence of indicated concentrations of each compound. Data are presented as mean values ± s.d. calculated across n = 2 replicates. **(D, E)** Western blot measurements **(D)** and their quantification **(E)** for p-SYK^Y525/526^ relative to total SYK in MeWo cells treated for 24 h with indicated concentrations of MTX-216, MTX-211, R406 and entospletinib. Antibodies were co-stained on the same blot. Data across different treatments conditions are normalized to DMSO-treated cells. **(F)** Differentially up-or down-regulated genes by MTX-216 and MTX-211 (relative to DMSO) following 24 h treatment of WM3918 and MeWo cells. Significant genes were identified as |log_2_(fold change)| ≥ 1 and adjusted *P* value (FDR) ≤ 0.05. **(G)** Number of significant genes up- or down-regulated by only one of the compounds or by both compounds in each of the WM3918 and MeWo cell lines. **(H)** Number of genes down-regulated specifically by MTX-216 in either or both of WM3918 and MeWo cell lines. **(I)** Top enriched kinase perturbation terms associated with 231 genes down-regulated specifically by MTX-216 (but not by MTX-211) in both WM3918 and MeWo cell lines.

We then asked whether the difference in the ability of MTX-216 and MTX-211 to inhibit SYK was seen in NF1^LoF^ cells. We exposed MeWo cells to MTX-216 and MTX-211 for 24 h and used Western blotting to measure dose-dependent changes in p-SYK^Y525/526^, autophosphorylation sites on SYK activation loop that are required for its kinase activity (40,41). As a control for SYK inhibition, we used two ATP-competitive SYK inhibitors, R406 and entospletinib (42,43). Consistent with its higher potency in inhibition of SYK in biochemical assays, MTX-216 reduced p-SYK^Y525/526^ more effectively in comparison with MTX-211 across both tested concentrations (1 and 3.2 μM) (Fig. 3D, E). MTX-216 at 1 μM inhibited p-SYK expression in MeWo cells by 50%, which was comparable to the effect of R406 and higher than the effect of entospletinib at the same concentration (Fig. E).

SYK function has been known primarily in the context of immune signaling in B cells, where it phosphorylates different proteins such as B-cell linker protein (BLNK) (17,44,45). We thus set out to test the effect of MTX-216 and MTX-211 (in comparison with R406 and entospletinib) on p-BLNK^Y96^ levels in IgM-stimulated Ramos cells, a B lymphocyte cell line. Western blot analysis showed that SYK inhibitors reduced p-BLNK levels in IgM-stimulated Ramos cells (Supplementary Fig. S3B, C). In addition, MTX-216 was more efficient in inhibition of p-BLNK in comparison with MTX-211.

To test whether the difference in inhibitory effects of MTX-216 and MTX-211 on SYK kinase was recapitulated at the level of transcription, we performed RNA sequencing on two NF1^LoF^ melanoma cell lines (MeWo and WM3918) exposed to either MTX-216 (at 1 μM), MTX-211 (at 1 μM), or vehicle (DMSO) for 24 h. Genes differentially expressed by at least two-fold by either MTX-216 or MTX-211 in each cell line were selected based on a statistical cutoff of FDR ≤ 0.05 (Fig. 3F, Supplementary Dataset S1). Among these genes, we focused on the subset differing in enrichment between the two compounds in both cell lines (Fig. 3G). Genes enriched specifically in MTX-216 treatment (but not in MTX-211 treatment) in both cell lines comprised of 231 down-regulated genes and 467 up-regulated genes (Fig. 3H, Supplementary Fig. S3D). We used these genes in Enrichr (30) and compared them statistically with gene expression signatures extracted from the Gene Expression Omnibus (GEO) database for kinase perturbation. The top kinase perturbation term was SYK drug inhibition (Fig. 3I, Supplementary Fig. S3E, and Supplementary Dataset S2), suggesting that MTX-216 at a concentration of 1 μM (more strongly than MTX-211) affects the expression of genes whose transcription is influenced by SYK inhibition.

Finally, we evaluated the effect of MTX-216 on its nominal targets and SYK activity *in vivo* by carrying out pharmacodynamic assessment of p-EGFR^Y1068^, p-AKT^S473^ and p-SYK^Y525/526^ in MTX-216 treated tumors (Supplementary Fig. 3F). Western blot analyses of WM3918 tumors excised from mice treated with a single dose of MTX-216 (at either 50 mg/kg or 100 mg/kg) revealed that this compound was comparatively more effective *in vivo* at reducing the expression of p-SYK compared to p-EGFR and p-AKT. Together, these data identify SYK as a target of MTX-216.

### Inhibition of SYK underlies increased sensitivity of NF1^LoF^ cells to MTX-216

To directly evaluate the importance of SYK for growth of NF1^LoF^ cells, we depleted the protein in two NF1^LoF^ melanoma cell lines (MeWo and YUHEF) using four specific siRNAs (targeting SYK exons 10, 11, 12 and 13) combined into a single pool. SYK knockdown led to reduced SYK protein levels in both cell lines (Fig. 4A). Following 3 and 6 days of inhibition by SYK siRNAs, we measured live cell counts and computed the normalized growth rate in SYK-depleted cells relative to cells treated with nontargeting (control) siRNA. We also compared the normalized growth rates in the presence of different doses of MTX-211, trametinib and pictilisib or vehicle (DMSO). SYK depletion in both MeWo and YUHEF led to a statistically significant decrease in their growth rates and enhanced their sensitivity to MTX-211, trametinib and pictilisib (Fig. 4B). To rule out the possibility of off-targeting by SYK siRNAs, we also examined the effects of each of the four SYK siRNAs independently. All individual siRNAs reduced SYK expression and growth rate in YUHEF cells (Supplementary Fig. S4A, B).

To test the impact of pharmacological inhibition of SYK on NF1^LoF^ melanoma cell growth, we treated three cell lines (MeWo, YUHEF, and WM3918) with the SYK inhibitor R406, individually or in combination with either MTX-211, trametinib or pictilisib. Inhibition of SYK by R406 reduced the growth rate in these cell lines by approximately 2-fold and enhanced their sensitivity to MTX-211, trametinib and pictilisib (Fig. 4C and Supplementary Fig. S4C). Together, these data show that both genetic and pharmacological inhibition of SYK significantly reduced the growth of NF1^LoF^ melanoma cells and increased their sensitivity to PI3K and MEK inhibitors. It is noteworthy, however, that the combination of SYK inhibition with MTX-211 did not exactly phenocopy the impact of MTX-216 alone. We speculate that this might be due to differences in mechanism of SYK inhibition by MTX-216 or possibly inhibition of other potentially significant targets by MTX-216.

Because treatment efficacy in NF1^LoF^ cells was correlated with an increase in the proportion of p-S6^Low^/Ki-67^Low^ subpopulation, we asked how p-S6 and Ki-67 levels would change in NF1^LoF^ cells following SYK inhibition. 72 h treatment with R406 reduced p-S6 and Ki-67 and increased the proportion of p-S6^Low^/Ki-67^Low^ subpopulation in both MeWo and YUHEF cells (Supplementary Fig. S4D). Because SYK inhibition enhanced the sensitivity of NF1^LoF^ cells to MEK inhibition, we also evaluated the effect of combined MEK and SYK inhibition. The combination of R406 with trametinib enhanced co-suppression of p-S6 and Ki-67 and increased the fraction of p-S6^Low^/Ki-67^Low^ subpopulation (Supplementary Fig. 4D).

### MTX-216 suppresses genes that regulate mitochondrial electron transport chain in NF1^LoF^ cells

To better understand the mechanisms underlying the superior efficacy of MTX-216 relative to other inhibitors, we extended our RNA sequencing analysis (described above) to treatments with additional inhibitors, including pictilisib (at 1 μM) and trametinib (at 0.1 μM). We also included a third NF1^LoF^ cell line (SKMEL113), which was treated for 24 h with either MTX-216 (at 1 μM), trametinib (at 0.1 μM), the combination of MTX-216 and trametinib (at 1 μM and 0.1 μM, respectively), or vehicle (DMSO). Gene set enrichment analysis (GSEA) of the effect of each individual compound on each cell line (relative to DMSO) confirmed the role of trametinib as an inhibitor of the ERK pathway, and pictilisib, MTX-216, and MTX-211 as inhibitors of the PI3K/AKT/mTOR pathway (Supplementary Fig. S5A and Supplementary Datasets S3, S4). To systematically compare gene expression changes across all 42 samples (including 3 cell lines, 6 distinct treatment conditions tested in 3 independent replicates), we used principal component analysis (PCA). PCA captured >80% of variance in the entire dataset by the first five principal components (Fig. 5A and Supplementary Datasets S5, S6). While the first two principal components (PC1 and PC2) associated mostly with cell line-specific differences, PC3 and PC4 were able to capture treatment-specific variations across the samples (Supplementary Fig. S5B). Changes induced by trametinib were revealed by a large shift in scores along PC4 for all three cell lines (Fig. 5B; purple arrows). MTX-216 treatment, on the other hand, led to a moderate shift along PC4, while inducing a strong shift along PC3 (Fig. 5B; blue arrows). The combination of trametinib and MTX-216 was additive, leading to substantial shifts in both PC4 and PC3 (Fig. 5B; red arrow), whereas pictilisib and MTX-211 induced only small changes along both principal components.

**Figure 4.**
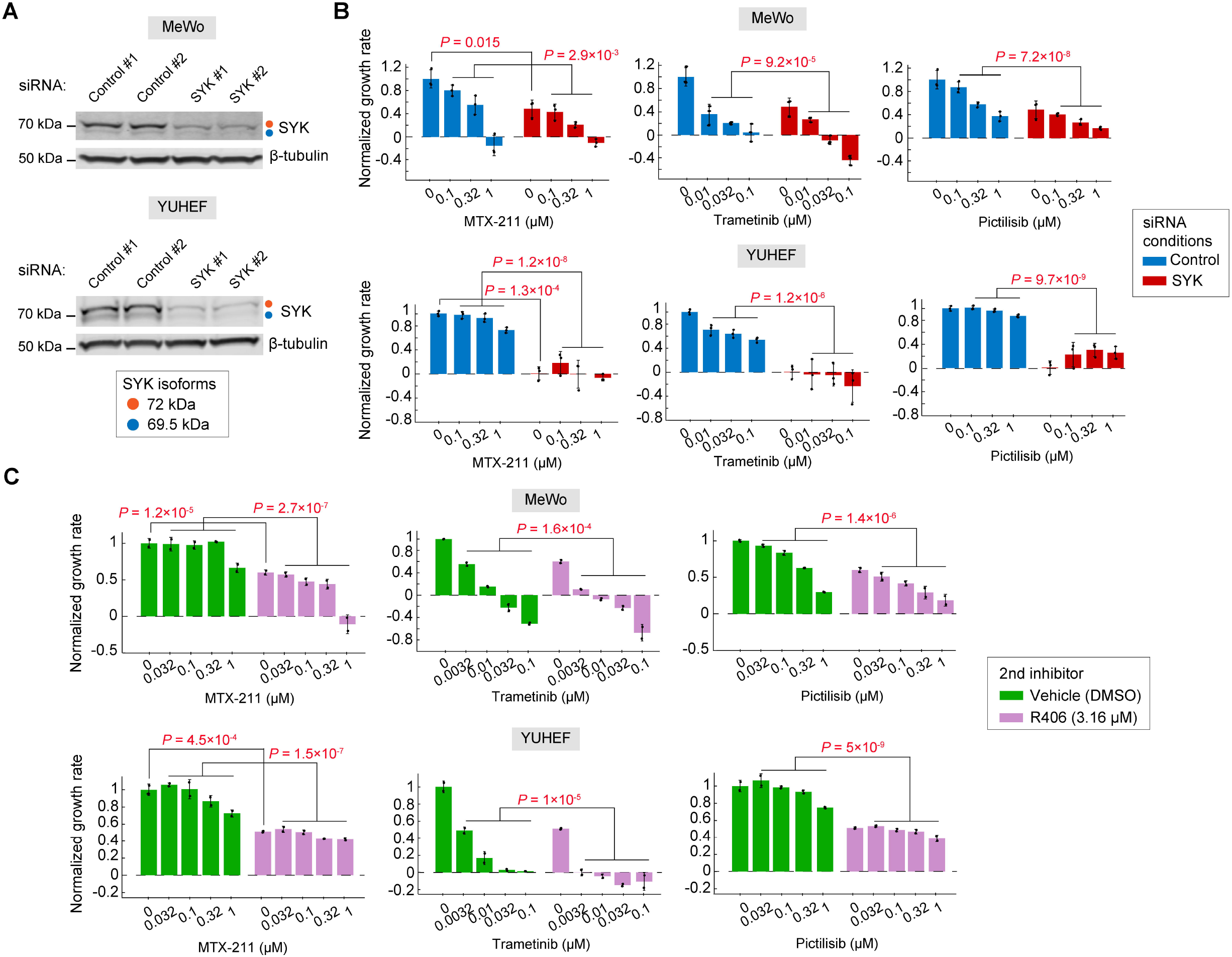
SYK inhibition or depletion reduces the growth of NF1^LoF^ melanoma cells and increases their sensitivity to both PI3K and MEK inhibitors. **(A)** SYK protein levels assessed by Western blots in MeWo (top) and YUHEF cells (bottom) following treatment with siRNAs targeting SYK isoforms or with non-targeting (control) siRNA for 72 h. Antibodies were co-stained on the same blot. Each siRNA experiment was performed in two independent replicates. **(B)** Normalized growth rates measured in MeWo (top) and YUHEF cells (bottom) following 72 h treatment with indicated concentrations of MTX-211, trametinib, or pictilisib (all dissolved in DMSO as vehicle) in the presence of siRNAs targeting SYK or non-targeting (control) siRNA (for 72 h). Data are presented as mean values ± s.d. calculated across n = 3 replicates. Statistical significance of the effect of SYK siRNA (versus control siRNA) in the absence of all inhibitors was determined by two-sided, two-sample *t*-test. Statistical significance of the effect of SYK siRNA (versus control siRNA) in the presence of inhibitors at three indicated concentrations was determined by two-way ANOVA. **(C)** Normalized growth rates measured in MeWo (top) and YUHEF cells (bottom) following 7-day treatments with indicated concentrations of MTX-211, trametinib, or pictilisib in combination with either R406 (at 3.16 μM) or vehicle (DMSO). Data are presented as mean values ± s.d. calculated across n = 2 replicates. Statistical significance of the effect of R406 (versus DMSO) in the absence of other inhibitors was determined by two-sided, two-sample *t*-test. Statistical significance of the effect of R406 (versus DMSO) in the presence of other inhibitors at four indicated concentrations was determined by two-way ANOVA.

**Figure 5.**
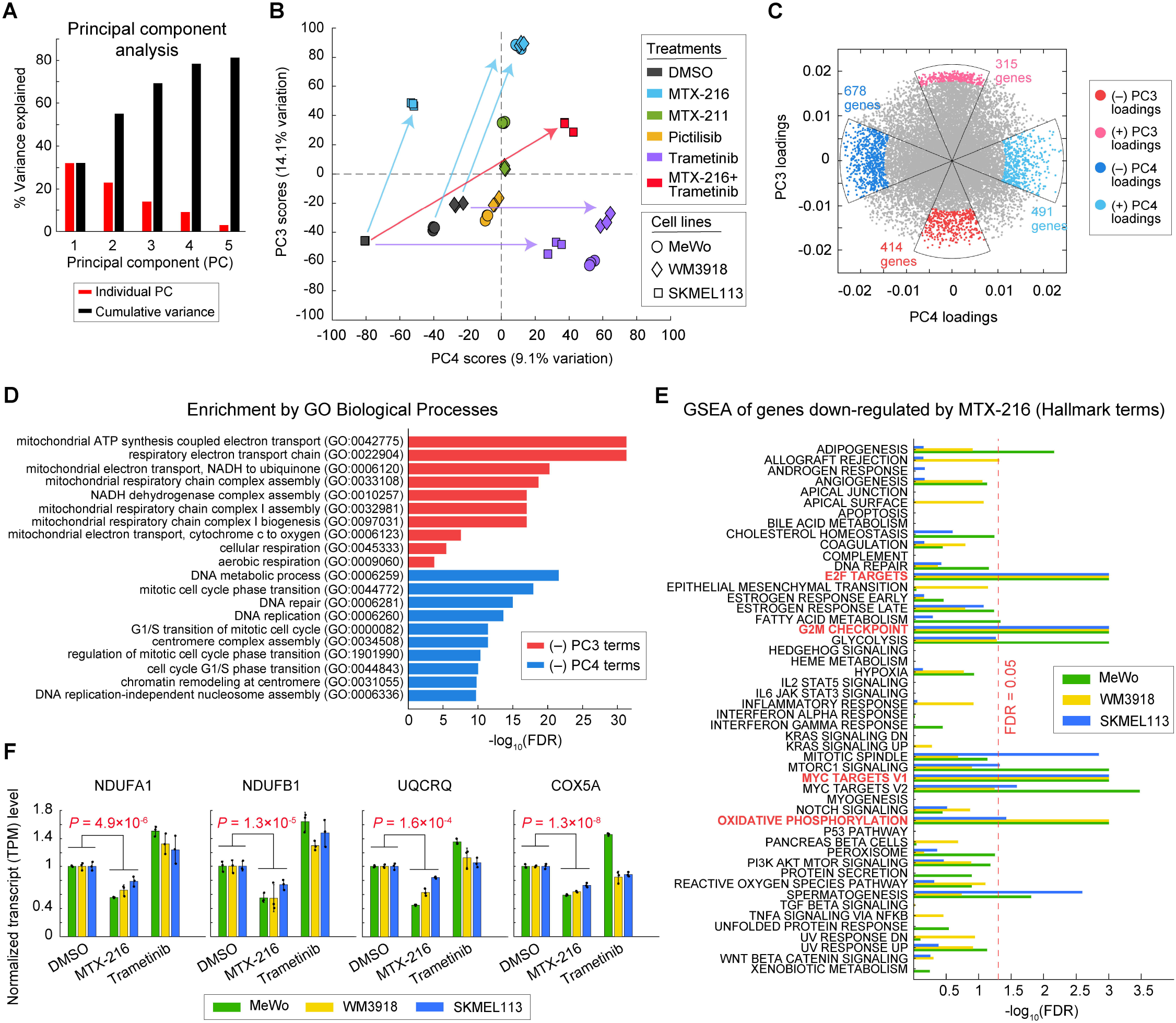
MTX-216 suppresses mitochondrial electron transport chain genes in NF1^LoF^ cells. **(A-C)** Principal component analysis of gene expression variations across 42 samples, representing 3 cell lines (MeWo, WM3918 and SKMEL113) and 24 h treatment conditions with MTX-216 (at 1 μM), MTX-211 (at 1 μM), pictilisib (at 1 μM), trametinib (at 0.1 μM), the combination of MTX-216 (at 1 μM) and trametinib (at 0.1 μM), or DMSO, each tested in 3 independent replicates. **(A)** Percentage of variance captured by the first five principal components (PCs). **(B)** PC3 and PC4 scores, capturing treatmentspecific variations in gene expression across the samples. Arrows show changes of scores (in PC3-PC4 space) by specific treatments relative to DMSO-treated cells. **(C)** PC3 and PC4 loadings, highlighting genes associated with distinct inhibitory effects on NF1^LoF^ cells. Gene loadings differentially enriched along either PC3 or PC4 are highlighted in color. **(D)** The top Gene Ontology (GO) terms with FDR ≤ 0.05 associated with significantly negative PC3 and significantly negative PC4 loadings. **(E)** Gene set enrichment analysis (GSEA) of genes down-regulated by MTX-216 in three cell lines (MeWo, WM3918 and SKMEL113) using consolidated Hallmark gene sets. Hallmark terms with FDR ≤ 0.05 in all three cell lines are highlighted in red. **(F)** DMSO-normalized gene expression (transcripts per million; TPM) levels of electron transport chain genes NDUFA1, NDUFB1, UQCRQ and COX5A in three cell lines following 24 h treatment with MTX-216 and trametinib. Data are presented as mean values ± s.d. calculated across n = 3 replicates. Statistical significance was determined by two-way ANOVA.

Because PC3 and PC4 separated the efficacious inhibitors independent of cell lines, we set out to use PC3 and PC4 loadings to identify genes associated with distinct inhibitory effects on NF1^LoF^ cells (Fig. 5C). The identified genes comprised 315 genes with strong positive PC3 loadings, 414 genes with strong negative PC3 loadings, 491 genes with strong positive PC4 loadings, and 678 genes with strong negative PC4 loadings. The top Gene Ontology (GO) terms associated with significantly negative PC4 loadings included processes involved in cell cycle progression, DNA replication and mitosis (Fig. 5D; blue bars, Supplementary Dataset S7). As expected from the PCA scores, these genes were suppressed profoundly in cells exposed to trametinib but were also down-regulated in cells treated with MTX-216 (Supplementary Fig. S5C). In contrast, genes associated with negative PC3 loadings were substantially down-regulated only in MTX-216 treated cells, but not in cells treated with other inhibitors (Supplementary Fig. S5C). The top GO terms associated with these genes included processes involved in cellular respiration and oxidative phosphorylation such as mitochondrial ATP synthesis and respiratory electron transport chain complex biogenesis and assembly (Fig. 5D; red bars, Supplementary Dataset S7). Consistent with these results, an independent GSEA using consolidated Hallmark gene sets (36) identified oxidative phosphorylation to be significantly associated with genes down-regulated by MTX-216 (Fig. 5E and Supplementary Datasets S1, S8, S9). Among the key genes affected were those that encode mitochondrial enzyme subunits involved in the respiratory electron transport chain, such as NADH:Ubiquinone Oxidoreductase Subunits A1 and B1(NDUFA1, NDUFB1), Ubiquinol-Cytochrome C Reductase Complex III Subunit VII (UQCRQ) and Cytochrome C Oxidase Subunit 5A (COX5A), which were all consistently down-regulated by MTX-216, but not by trametinib, in all three cell lines (Fig. 5F). Proliferation hallmarks (such as E2F targets, MYC targets and G2/M checkpoint), however, were down-regulated by both trametinib and MTX-216 (Fig. 5E, Supplementary Fig. S5D). Together, these data show that while both MTX-216 and trametinib are effective in inhibition of cell proliferation in NF1^LoF^ melanoma cells, the two compounds are distinguishable with respect to their effect on the suppression of genes that regulate mitochondrial electron transport chain.

### Mitochondrial respiratory chain is associated with poor survival in NF1^LoF^ melanomas patients and inhibitable by MTX-216 or SYK inhibition

To test the clinical relevance of SYK kinase and genes encoding mitochondrial electron transport chain (ETC), we assessed their association with NF1 mutation and patient phenotype across melanoma tumors in The Cancer Genome Atlas (TCGA) database. For this purpose, we first grouped patient-derived melanoma tumors based on their NF1 mutation status. Patients bearing “truncating” driver mutations were classified as NF1^LoF^ (39 patients); patients bearing “missense” or “splice” driver mutations were classified as NF1^Mutant^ (9 patients); and patients with no NF1 mutations or only “passenger” NF1 mutations were classified as NF1^Other^ (421 patients). We then analyzed the frequency of SYK mutations across each group of patients. Interestingly, the NF1^LoF^ and NF1^Mutant^ groups showed higher numbers of SYK mutations (including mostly missense mutations) compared to the NF1^Other^ group (Supplementary Fig. S6A). From these mutations, however, we were not able to identify any obvious patterns of altered SYK activity. Therefore, we extended our analysis to the gene expression data and asked whether there was any difference in SYK pathway activity between different groups of patients. As a surrogate for SYK activity, we used the SYK drug inhibition gene sets, including genes whose expression changes following treatment with SYK inhibitor (Supplementary Dataset S4). Gene set enrichment analysis showed a slight increase in SYK activity in the NF1^LoF^ and NF1^Mutant^ groups relative to the NF1^Other^ group based on the normalized enrichment score (NES), but the difference was not statistically significant (Supplementary Fig. S6B). We also did not observe any significant differences in the overall expression of ETC genes (Respiratory Electron Transport Chain GO term; GO:0022904, Supplementary Dataset S4) in tumors across three groups of melanoma patients (Supplementary Fig. S6C). However, when we focused our analysis on each group separately and asked whether the expression of ETC genes correlated with patient survival, we found that within the NF1^LoF^ group (or NF1^LoF^ and NF1^Mutant^ groups combined), patients with ETC^High^ tumors had a significantly shorter overall survival compared to the patients with ETC^Low^ tumors (Figure 6a; left and middle panels). Within the NF1^Other^ cohort of patients, however, there was no significant difference between the overall survival and whether tumors belonged to ETC^High^ or ETC^Low^ subgroups (Figure 6a; right panel). To test whether such association within the NF1^LoF^/NF1^Mutant^ groups might be due to confounding health risks, we assessed the patients’ diagnosis stage, but we did not observe any significant differences among NF1^LoF^/NF1^Mutant^ patients with ETC^High^ tumors versus those with ETC^Low^ tumors (Supplementary Fig. S6D). These data suggest that melanoma tumors with NF1 mutations may rely on mitochondrial respiratory chain for progression.

**Figure 6.**
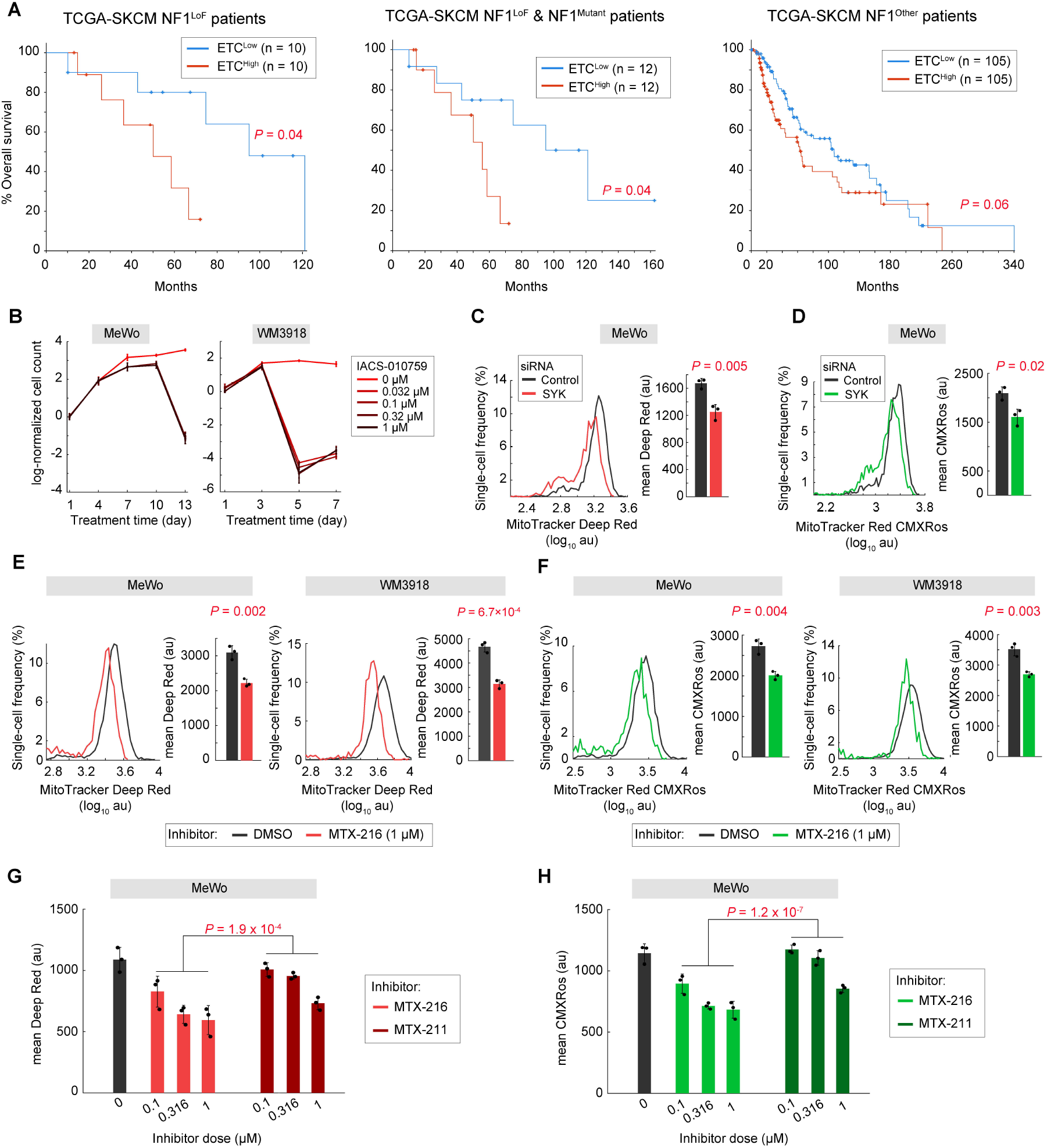
Mitochondrial respiratory chain is associated with poor survival in NF1^LoF^ melanomas patients and inhibitable by MTX-216 or SYK inhibition. **(A)** Overall survival analysis of patients with ETC^High^ or ETC^Low^ tumors. The analysis was performed in three groups of patients: patients with NF1^LoF^ tumors (left), patients with either NF1^LoF^ or NF1^Mutant^ tumors (middle), and patients with no significant NF1 mutations; NF1^Other^ (right). Statistical significance was determined by two-sided logrank (Mantel-Cox) test. **(B)** Log-normalized changes in live cell count following the exposure of two cell lines MeWo and WM3918 to indicated doses of IACS-010759 for a period of 13 and 7 days, respectively. Data are presented as mean values ± s.d. calculated across n = 2 replicates. **(C, D)** Changes in mitochondrial mass measured by MitoTracker Deep Red **(C)**, and mitochondrial membrane potential measured by MitoTracker Red CMXRos **(D)**, in MeWo cells in the presence of SYK siRNAs or nontargeting (control) siRNA for 72 h. Single-cell intensities (left) and the population-averaged data (right) are shown for each measurement. Population-averaged data are presented as mean values ± s.d. calculated across n = 3 replicates. Statistical significance was determined by two-sided, two-sample *t*-test. **(E, F)** Changes in mitochondrial mass measured by MitoTracker Deep Red **(E)**, and mitochondrial membrane potential measured by MitoTracker Red CMXRos **(F)**, in MeWo and WM3918 cells following 48 h treatment with MTX-216 (at 1 μM) or DMSO. Single-cell intensities (left) and the population-averaged data (right) are shown for each measurement. Population-averaged data are presented as mean values ± s.d. calculated across n = 3 replicates. Statistical significance was determined by two-sided, two-sample *t*-test. **(G, H)** Changes in mitochondrial mass measured by MitoTracker Deep Red **(G)**, and mitochondrial membrane potential measured by MitoTracker Red CMXRos **(H)**, in MeWo cells following 48 h treatment with different concentrations of MTX-216, MTX-211, or DMSO. Population-averaged data are presented as mean values ± s.d. calculated across n = 3 replicates. Statistical significance was determined by two-way ANOVA.

To directly evaluate the importance of mitochondrial respiratory chain for growth of NF1^LoF^ cells, we used IACS-010759, a clinical-grade small molecule inhibitor of complex I of the mitochondrial electron transport chain. IACS-010759 inhibited growth in a dose-dependent fashion in all tested cell lines (Fig. 6B and Supplementary Fig. S6E). Using the area over of the dose-response curve to compare overall drug sensitivity, we found the NF1^LoF^ cell lines to be similarly sensitive to both IACS-010759 and MTX-216, and their sensitivities to these compounds were greater than their sensitivity to R406 (Supplementary Fig. S6E, F). When combined with trametinib, however, MTX-216 led to the largest reduction in growth rate (Supplementary Fig. S6E, F). Together, these data support the hypothesis that NF1^LoF^ melanoma cells depend on mitochondrial respiratory chain for growth, and that MTX-216 and trametinib target orthogonal vulnerabilities, leading to enhanced benefit when used in combination.

Because genes down-regulated by MTX-216 were involved in the mitochondrial respiratory chain, we asked whether this effect was due to a global reduction in mitochondrial mass or function because of SYK inhibition by this compound. We thus used the fluorescent probes, MitoTracker Deep Red and MitoTracker Red CMXRos, to quantitate the mitochondrial mass and membrane potential in NF1^LoF^ cells, respectively (46). Measurements were performed by fluorescence microscopy in two cell lines (MeWo and WM3918) following treatment with either SYK siRNA or a non-targeting control siRNA for 72 h, or with MTX-216 (at 1 μM) or vehicle (DMSO) for 48 h. Both SYK knockdown (Fig. 6C, D) and MTX-216 treatment (Fig. 6E, F) shifted cell populations towards lower levels of both mitochondrial markers. To compare the effect of MTX-216 and MTX-211 on mitochondria, we performed the assays in MeWo cells following 48 h treatment with increasing concentrations of either compound. MTX-216 was significantly more potent than MTX-211 in reducing both mitochondrial markers (Fig. 6G, H and Supplementary Fig. S7). Together, these results are consistent with RNA sequencing data, suggesting that the efficacy of MTX-216 through SYK inhibition is mediated by the suppression of mitochondrial mass and electron transport chain function.

## Discussion

We performed a targeted kinase inhibitor screen to identify vulnerabilities whose inhibition, individually or in combination with MEK/ERK or PI3K/mTOR inhibitors, would be effective in blocking NF1^LoF^ melanoma cells. We found that the efficacy of kinase inhibitors was determined by their ability to co-inhibit the proliferation marker Ki-67 and the ribosomal S6 phosphorylation (p-S6). While MEK/ERK inhibitors and PI3K inhibitors led to heterogeneous responses, we identified a tool compound, MTX-216, that effectively suppressed both Ki-67 and p-S6 in NF1^LoF^ melanoma cells, inducing efficacious anti-tumor responses in these cells. MTX-216 was originally developed as a dual PI3K/EGFR inhibitor. However, systematic analysis showed that its high efficacy in NF1^LoF^ cells depended on its ability to coinhibit SYK kinase. Therefore, the study of MTX-216 served as a tool to identify SYK as a new vulnerability which when exploited, especially in combination with MEK inhibition, can lead to efficacious responses in NF1^LoF^ melanoma cells. The greater efficacy of MTX-216 in NF1^LoF^ cells, in comparison with BRAF^V600E^ cells, is consistent with this compound being able to inhibit SYK, a nonreceptor tyrosine kinase that can be activated by multiple upstream receptors and initiates downstream signaling through RAS (15,40,47–49). Nevertheless, our data prompt additional future investigations to identify the role of SYK and its associated adaptors in relationship to RAS signaling and tumorigenesis in NF1^LoF^ melanomas. Such studies may provide a path to exploit SYK dependency to selectively block NF1^LoF^ melanoma tumors.

The discovery of SYK as a vulnerability in NF1^LoF^ melanomas based on the high efficacy of a tool compound (MTX-216) in these cells represents an unconventional approach that leverages mechanistic, poly-pharmacology drug studies to identify novel cancer targets (14). There are many instances of kinase inhibitors, whose anti-tumor efficacy was found to be due to their ability to inhibit kinases other than their originally intended targets (50). This is not surprising because all kinases bind to a common substrate (i.e., ATP), which underlies their great potential for promiscuity. Therefore, individual kinase inhibitors may lead to unpredicted activity in the combination setting, resulting from unique biological activities that could be exploited to discover new mechanisms as well as novel actionable dependencies in cancer cells.

Our analysis provides insight into how MTX-216 promotes anti-tumor effects in NF1^LoF^ melanoma cells by suppressing a group of genes that regulate mitochondrial electron transport chain and function. The expression of these genes is associated with poor survival in patients with NF1^LoF^ melanomas. Therefore, MTX-216 may block NF1^LoF^ melanoma cells by targeting a mitochondrial vulnerability associated with a specific SYK-dependent signaling context that is required for NF1^LoF^ tumor progression. This is consistent with the sensitivity of NF1^LoF^ melanoma cells to pharmacological inhibition of mitochondrial electron transport chain either by IACS-010759 as demonstrated in this study, or by phenformin as reported previously (10).

The role of SYK kinase in regulation of diverse pathways involved in cellular homeostasis, including mitochondrial function and oxidative metabolism, has been already well documented. In B cells, for example, SYK is activated by the B cell receptor pathway, regulating multiple signaling pathways involved in differentiation and proliferation, as well as oxidative phosphorylation and mitochondrial oxidation of fatty acids (15,19,45). Blocking oxidative metabolism via SYK inhibition has been shown to induce anti-tumor effects in acute myeloid leukemia, B cell chronic lymphocytic leukemia, and diffuse large B cell lymphoma (17–20). Furthermore, SYK activation has been shown to occur, at least in part, in proximity to the respiratory chain, where a pool of SYK localizes to the mitochondrial intermembrane space, controlling cellular response to oxidative stress in diverse cell types (16). Future studies may leverage these findings to elucidate the mechanistic connection between the SYK pathway activity and mitochondrial oxidative metabolism in NF1^LoF^ melanomas.

## Authors’ Contributions

C.A., C.W., E.Z., D.G.B. and C.F.-M. generated reagents, performed the experiments and analyzed the experimental data. C.A. performed bioinformatics and statistical analyses. C.A. and M.F.-S. wrote the manuscript. All authors reviewed and assisted with manuscript preparation and writing. M.F.-S. and J.S.-L. supervised the work.

## Acknowledgments

We thank the University of Virginia Advanced Microscopy Facility, University of Virginia Flow Cytometry Core Facility, University of Michigan Advanced Genomics Core, Kevin Janes, David Kashatus, Jennifer Kashatus, Armand Bankhead, and members of the Fallahi-Sichani and Sebolt-Leopold laboratory for help and discussion. This work was supported by the Turner-McConnell Fund for Drug Discovery and the University of Michigan Rogel Cancer Center Support Grant (P30-CA046592), University of Virginia Cancer Center Support Grant (P30-CA044579), Department of Defense PRCRP Career Development Award W81XWH1810427 (to MFS), NIH grants U54-CA274499, R35-GM133404 and R01-CA249229 (to MFS), NIH grants R01-CA220199 and R01-CA241764 (to JSL), and the University of Michigan Rackham Merit Fellowship (to CA).

