## Supplementary Figures for "Systematic analysis uncovers SYK dependency in NF1^LoF^ melanoma cells"

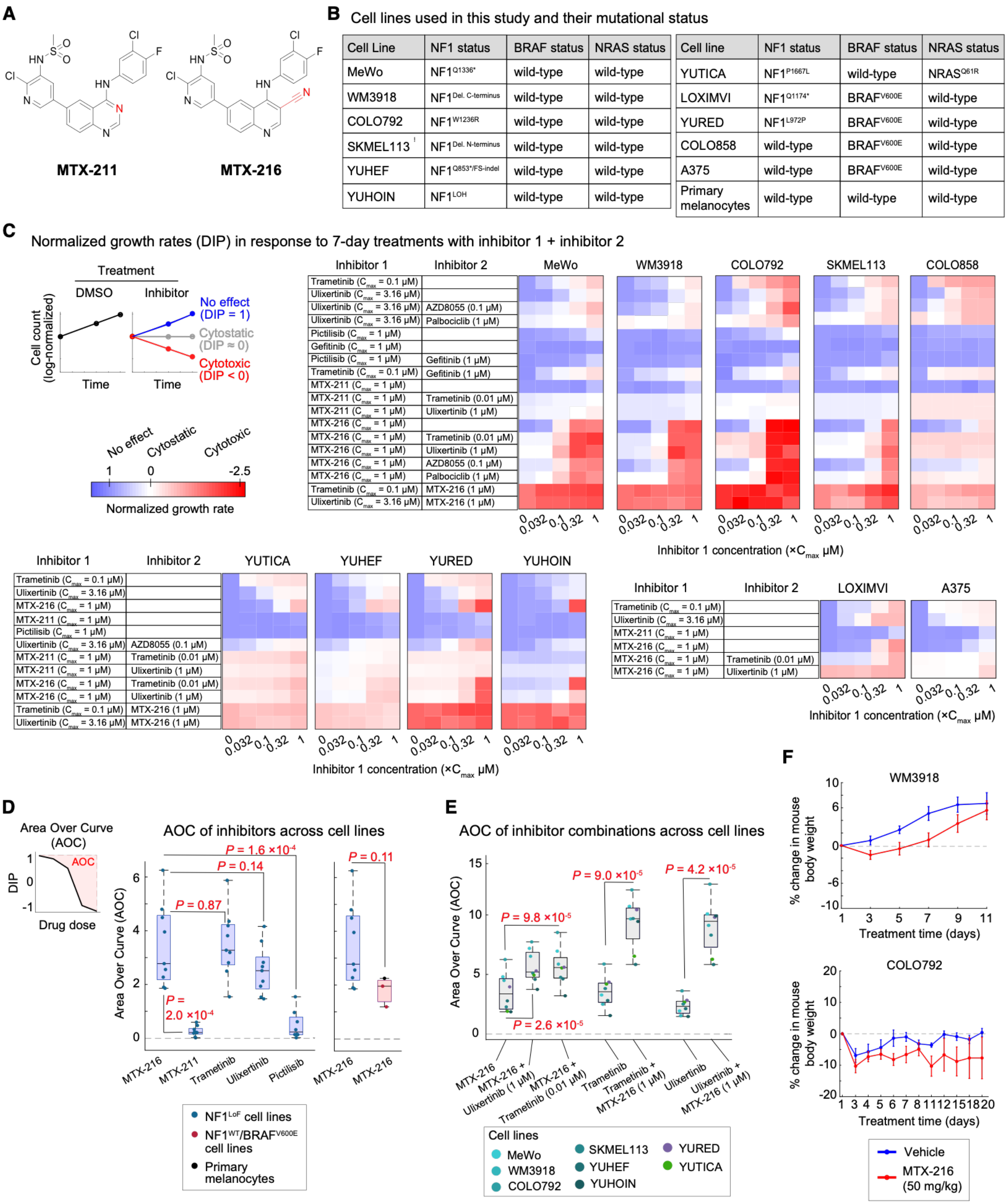


**Supplementary Figure S1. A targeted compound screen identifies MTX-216 as an efficacious inhibitor of NF1$^{LoF}$ melanoma cells. (A)** The chemical structures of MTX-216 (N-(2-chloro-5-(4-((3-chloro-4-fluorophenyl)amino)-3-cyanoquinolin-6-yl)pyridin-3-yl)methanesulfonamide) and MTX-211

(N-(2-chloro-5-(4-((3-chloro-4-fluorophenyl)amino)quinazolin-6-yl)pyridin-3-yl)methanesulfonamide). The key difference between the two compounds is the substitution at the 3-position of the ring system highlighted in red. The nitrogen at the 3-position of the quinazoline core structure of MTX-211 was replaced with a carbon and 3-cynano group to yield the quinoline core structure of MTX-216. **(B)** List of cell lines used in this study and their NF1, BRAF, and NRAS mutation status. **(C)** Dose-dependent changes in normalized growth rates (a.k.a. DIP rates) calculated by dividing the 7-day average net growth rate for inhibitor-treated cells to that measured for vehicle (DMSO)-treated cells across cell lines with different NF1/BRAF/NRAS mutation status. The average net growth rates for each condition were calculated from measurements of live cell count (across two replicates) at four timepoints (including 1, 3, 5, and 7 days). Normalized growth rates < 0 indicate a net cell loss (i.e., inhibitor-induced cytotoxicity), a value of 0 represent no change in viable cell number (i.e., cytostasis), a value > 0 indicates a net cell gain, and a value of 1 represents no effect as cells grow in the presence of inhibitor at the same rate as in the DMSO condition. For combination, the concentration of one inhibitor was varied between 0 and the indicated $C_{max}$ value in the presence of a fixed concentration of a second inhibitor. **(D, E)** Area over the dose-response curve (AOC) for different inhibitors **(D)** and their combinations **(E)** across $NF1^{LoF}$ and $NF1^{WT}$ melanoma cell lines. AOC value of 0 indicates no drug response. Statistical significance was determined by two-sided, paired-sample *t* test when comparison was made between the same group of cell lines (left plot in **D** and **E**) and two-sided, two-sample *t* test when comparison was made between different groups of cell lines (right plot in **D**). **(F)** Body weight change in mice implanted subcutaneously with WM3918 and COLO792 tumor cells following intraperitoneal (IP) treatments with MTX-216 or vehicle in two different dosing regimens as described in Methods. Data represent mean values ± s.e.m across 4 mice per group (in case of COLO792) or 5 mice per group (in case of WM3918).

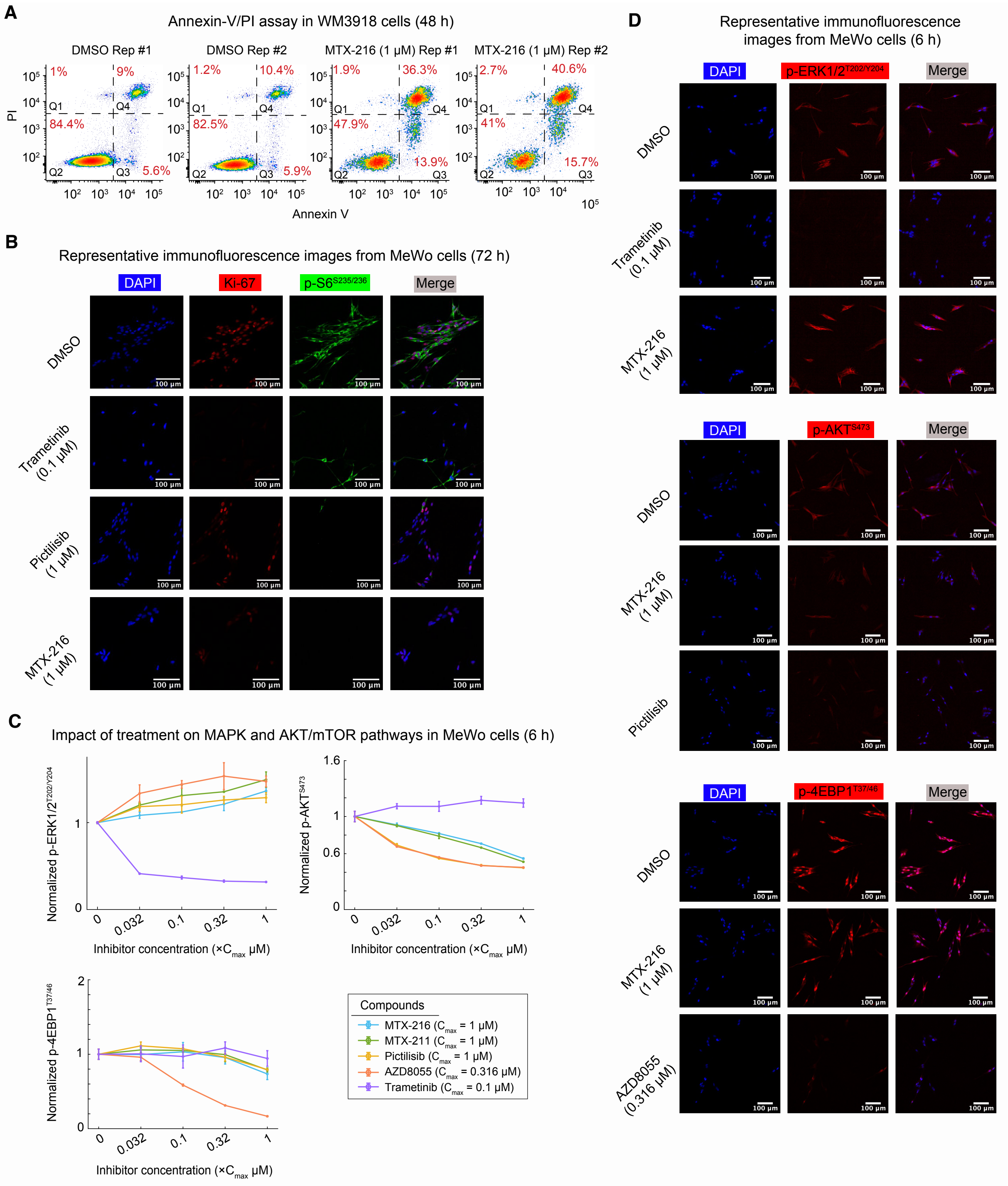


**Supplementary Figure S2. MTX-216 co-suppresses Ki-67 and p-S6 in NF1$^{LoF}$ cells. (A)** Annexin V and propidium iodide (PI) stained WM3918 cells treated with DMSO or MTX-216 (1 μM) for 48 h. Annexin V$^{-}$/PI$^{-}$ indicate living cells (Q2). Annexin V$^{+}$/PI$^{-}$ cells indicate cells undergoing early apoptosis (Q3). Annexin V$^{+}$/PI$^{+}$ indicate late apoptotic/necrotic cells (Q4). Annexin V$^{-}$/PI$^{+}$ indicate mechanically damaged cells (Q1). Percentages of cells in each quadrant are shown in red. **(B)** Representative multiplexed immunofluorescent images of Ki-67 and phosphorylated S6 ribosomal protein (p-$S6^{S235/S236}$), co-stained in MeWo cells following 72 h treatment with indicated concentrations of trametinib, pictilisib, MTX-216 or DMSO. Each experiment was repeated twice independently with similar results. Scale bars represent 100 µm. **(C)** Normalized dose-response changes in p-$ERK1/2^{T202/Y204}$**,** p-$AKT^{S473}$, and p-$4EBP1^{T37/46}$ in drug treated MeWo cells (6 h). Data are presented as mean values ± s.d. calculated across n = 2 replicates. Data across different treatments conditions are normalized to DMSO-treated cells. **(D)** Representative multiplexed immunofluorescent images of phosphorylated ERK (p-$ERK1/2^{T202/Y204}$), phosphorylated AKT (p-$AKT^{S473}$), and phosphorylated 4EBP1 (p-$4EBP1^{T37/46}$) in MeWo cells following 6 h treatment with indicated concentrations of trametinib, MTX-216, AZD8055, or DMSO. Each experiment was repeated twice independently with similar results. Scale bars represent 100 µm.

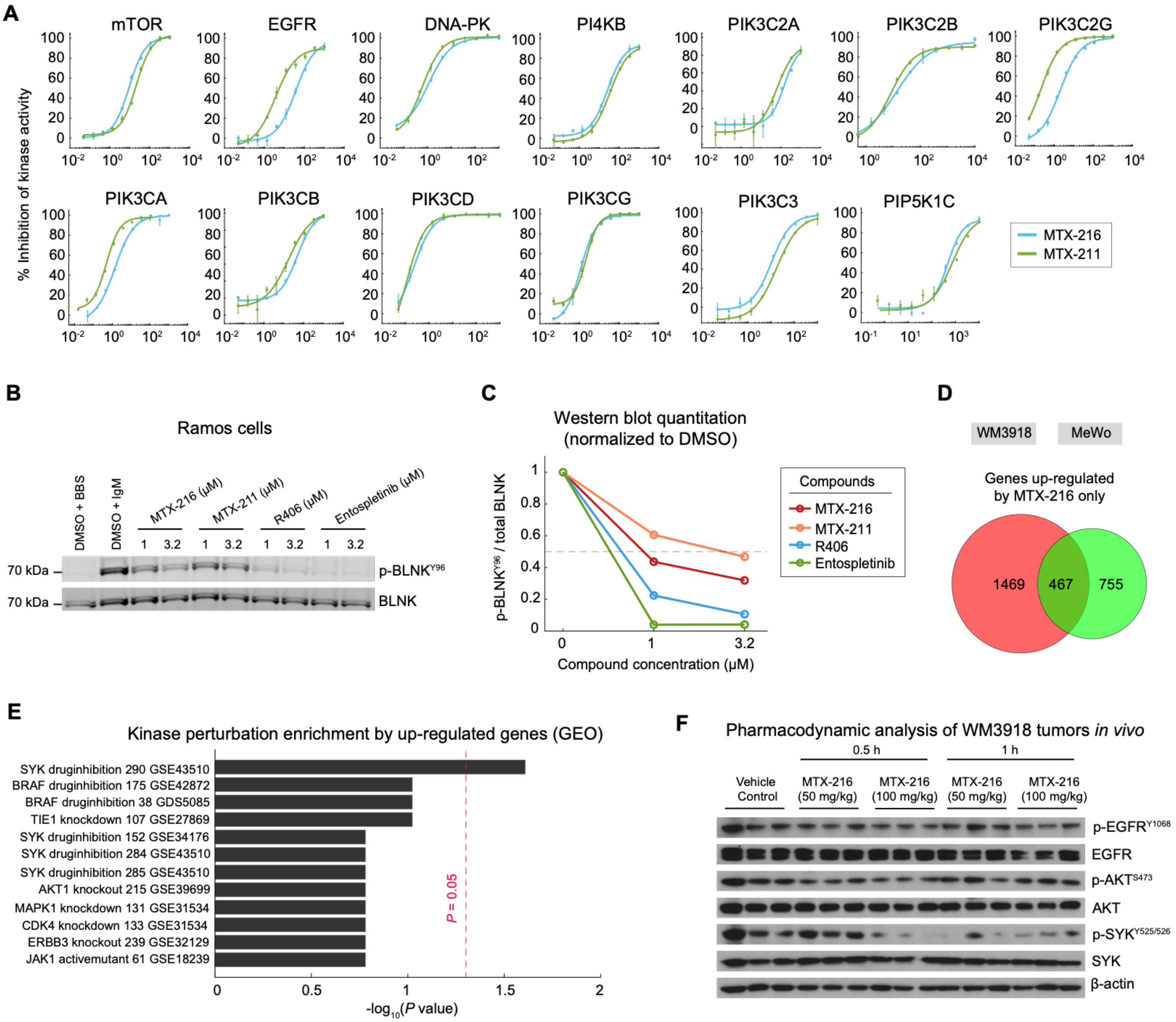


**Supplementary Figure S3. SYK kinase is inhibited by MTX-216 *in vitro* and *in vivo*. (A)** Dose-dependent inhibition of kinase activities by MTX-216 and MTX-211. Data are presented as mean values ± s.d. calculated across n = 2 replicates. Kinase inhibition assays (using Z'-LYTE and Adapta platforms) were performed in the presence of ATP at a concentration of $K_{m,app}$ for each target. **(B, C)** Western blot measurements **(B)** and their quantification **(C)** for p-BLNK$^{Y96}$ relative to total BLNK in Ramos cells treated for 6 h with indicated concentrations of MTX-216, MTX-211, R406 and entospletinib. Cells were stimulated with 25 µg/mL IgM or BBS for 3 min after drug treatment. Antibodies were co-stained on the same blot. Data across different treatments conditions are normalized to (DMSO + IgM)-treated cells. **(D)** Number of genes up-regulated specifically by MTX-216 in either or both of WM3918 and MeWo cell lines. **(E)** Top enriched kinase perturbation terms associated with 467 genes up-regulated specifically by MTX-216 (but not by MTX-211) in both WM3918 and MeWo cell lines. **(F)** SYK, EGFR, AKT total protein and phosphorylated protein levels assessed by Western blots in excised WM3918 xenograft tumors following intraperitoneal injection (IP) treatment with MTX-216 (50 mg/kg or 100 mg/kg) for 0.5 or 1 h. Experiments were performed in 3 mice per condition. Antibodies were stained on separate blots.

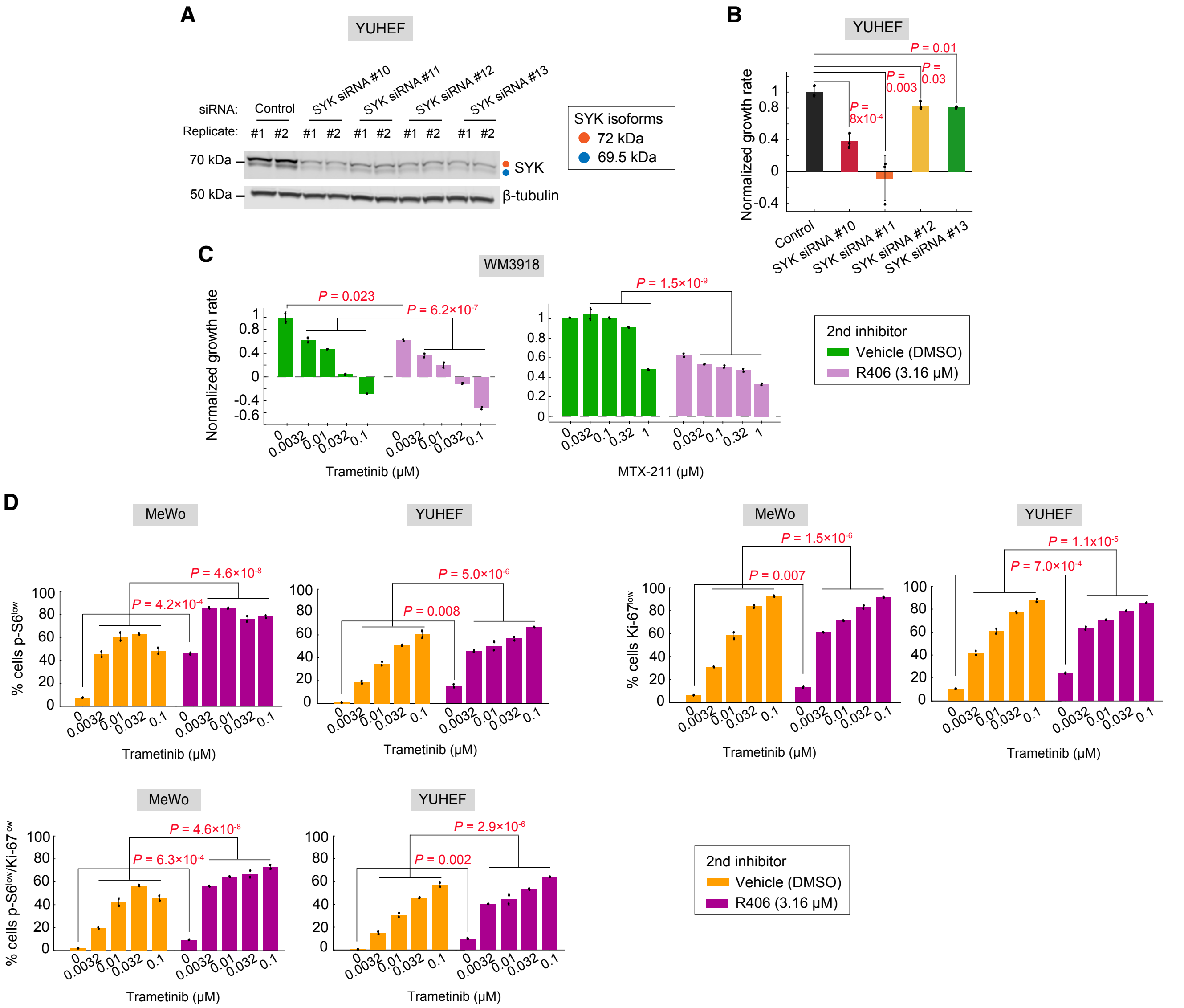

**Supplementary Figure S4. SYK inhibition reduces the growth of NF1$^{LoF}$ melanoma cells and increases their sensitivity to PI3K and MEK inhibition. (A)** SYK protein levels assessed by Western blots in YUHEF cells following treatment with individual SYK siRNAs or with non-targeting (control) siRNA for 72 h. siRNA numbers correspond to the last two digits of the Dharmacon catalogue numbers (e.g., J-003176-12 = SYK siRNA #12). Each siRNA experiment was performed in two independent replicates. **(B)** Normalized growth rates measured in YUHEF cells following treatment with indicated SYK siRNAs or non-targeting (control) siRNA (72 h). Data are presented as mean values ± s.d. calculated across n = 3 replicates. Statistical significance of the effect of each SYK siRNA (versus control siRNA) was determined by two-sided, two-sample *t*-test. **(C)** Normalized growth rates measured in WM3918 cells following 7-day treatments with indicated concentrations of MTX-211 or trametinib (both dissolved in DMSO as vehicle) in combination with either R406 (at 3.16 µM) or vehicle (DMSO). Data are presented as mean values ± s.d. calculated across n = 2 replicates. Statistical significance of the effect of R406 (versus DMSO) in the absence of the other inhibitors was determined by two-sided, two-sample t-test. Statistical significance of the effect of R406 (versus DMSO) in the presence of other inhibitors at four indicated concentrations was determined by two-way ANOVA. **(D)** Percentage of cells that were p-S6$^{low}$, Ki-67$^{low}$, or p-S6$^{low}$/Ki-67$^{low}$ in MeWo or YUHEF cells following 72 h treatment with indicated concentrations of trametinib in combination with either R406 (at 3.16 μM) or vehicle (DMSO). Data are presented as mean values ± s.d. calculated across n = 2 replicates. Statistical significance of the effect of R406 (versus DMSO) was determined by two-sided, two-sample t-test. Statistical significance of the effect of R406 (versus DMSO) in the presence of trametinib at four indicated concentrations was determined by two-way ANOVA.

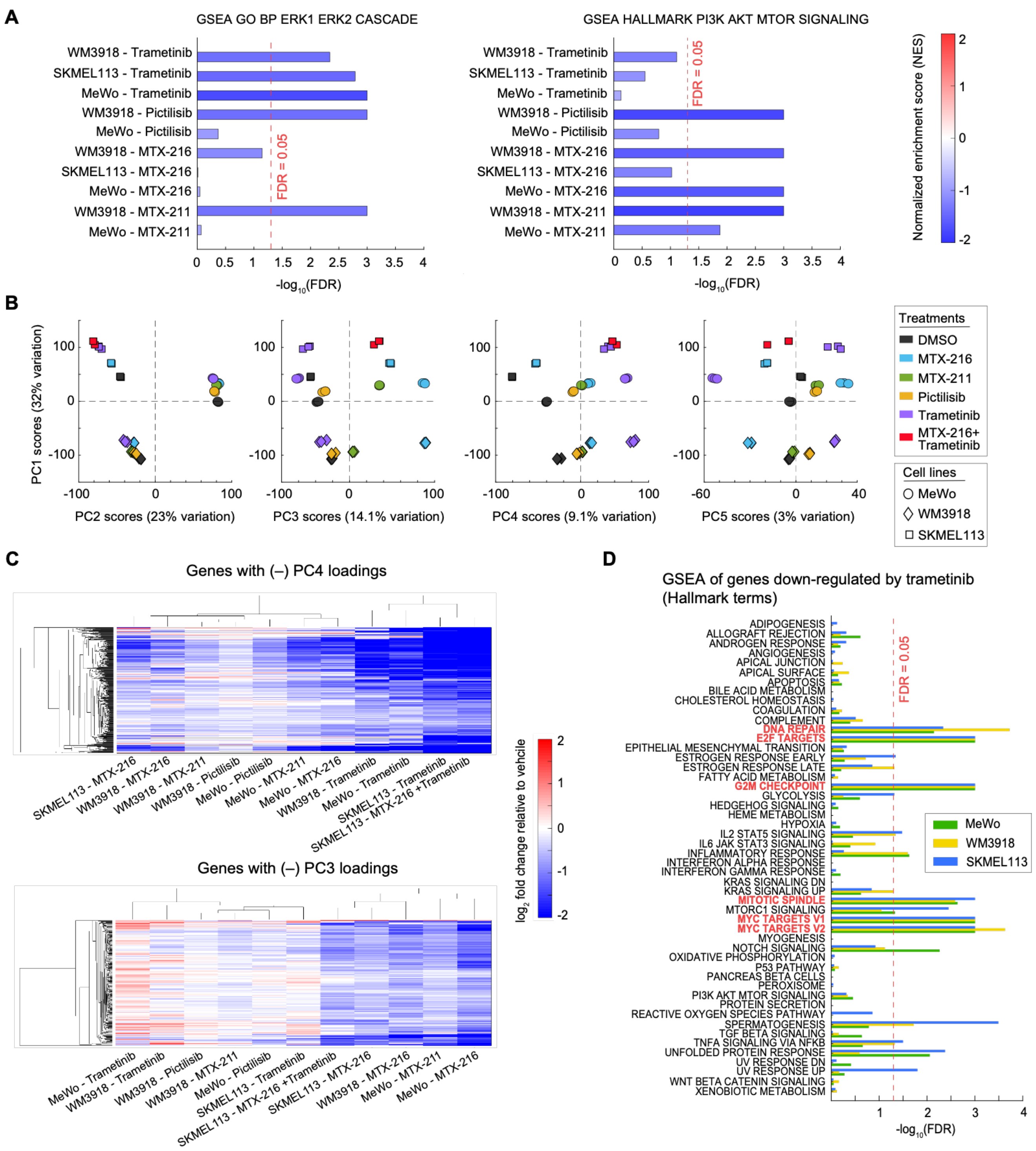


**Supplementary Figure S5. Statistical and bioinformatics analyses identify genes differentially enriched by trametinib versus MTX-216 in NF1$^{LoF}$ cells. (A)** Gene set enrichment analysis (GSEA) in three cell lines (MeWo, WM3918 and SKMEL113) treated with trametinib, pictilisib, MTX-211, or MTX-216 using GO BP ERK1 ERK2 Cascade and Hallmark PI3K AKT MTOR Signaling gene sets. **(B)** Principal component analysis of gene expression variations across 42 samples, representing 3 cell lines (MeWo, WM3918 and SKMEL113) and 24 h treatment conditions with MTX-216 (at 1 μM),

MTX-211 (at 1 μM), pictilisib (at 1 μM), trametinib (at 0.1 μM), the combination of MTX-216 (at 1 μM) and trametinib (at 0.1 μM), or DMSO, each tested in 3 independent replicates. PC1 and PC2 scores capture cell line-specific variations, while PC3 and PC4 scores capture inhibitor-specific variations in gene expression across the samples. **(C)** Hierarchical clustering of $\log_2$ fold changes (relative to DMSO) in the expression of genes with significantly negative PC4 loadings (top) or genes with significantly negative PC3 loadings (bottom) following treatment with indicated inhibitors. **(D)** Gene set enrichment analysis (GSEA) of genes down-regulated by trametinib in three cell lines (MeWo, WM3918 and SKMEL113) using consolidated Hallmark gene sets. Hallmark terms with FDR $\leq 0.05$ in all three cell lines are highlighted in red.

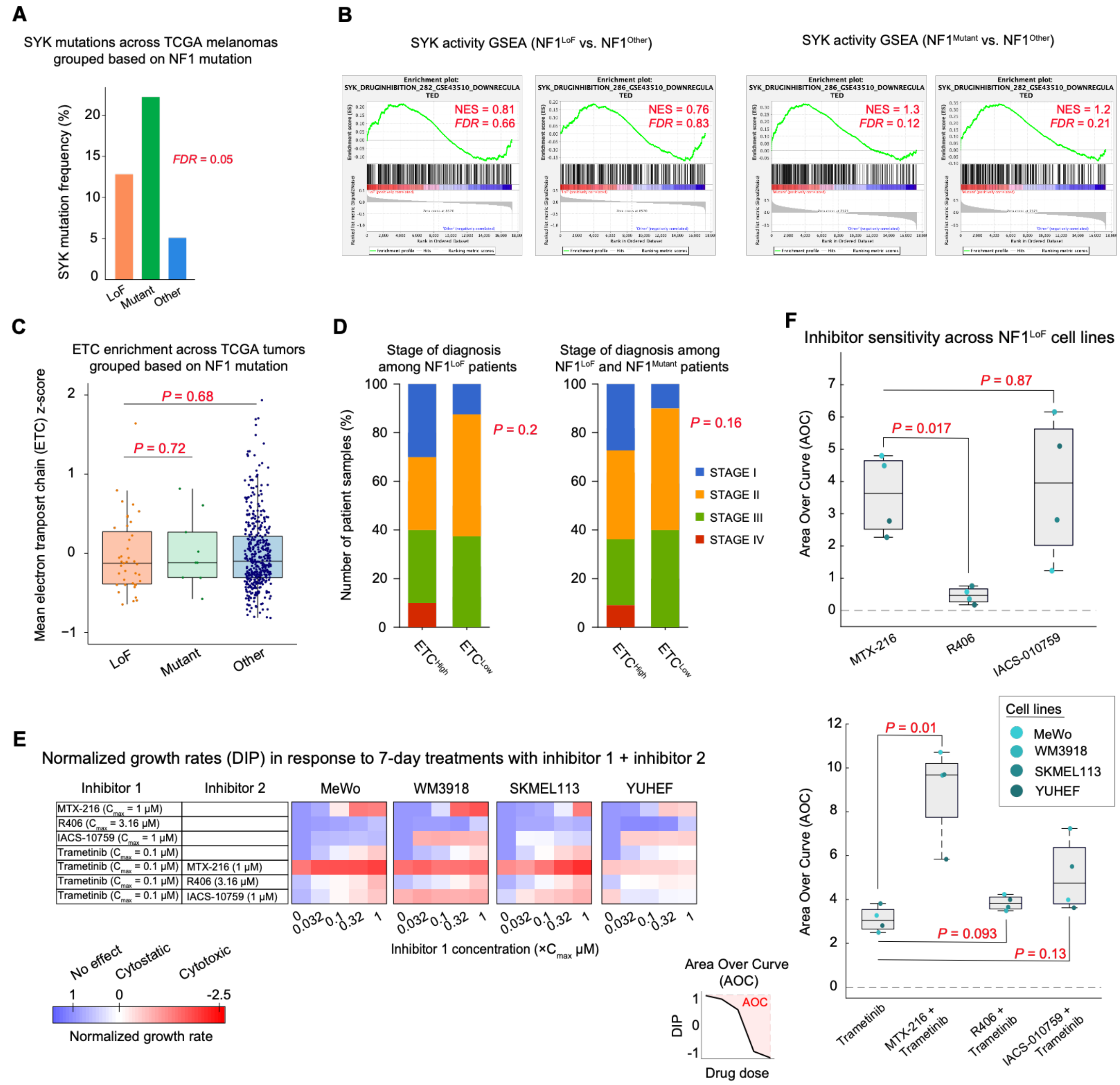


**Supplementary Figure S6. TCGA-SKCM patient analysis of SYK and electron transport chain and IACS-010759 drug response in NF1$^{LoF}$ melanoma cells. (A)** SYK mutation frequency across NF1 mutation groups in TCGA-SKCM patients. Statistical significance was determined with Chi-squared test followed by Benjamini-Hochberg FDR correction. **(B)** Gene set enrichment analysis (GSEA) plots comparing NF1$^{LoF}$ versus NF1$^{Other}$ patients or NF1$^{Mutant}$ versus NF1$^{Other}$ patients in terms of SYK drug inhibition gene signatures. Normalized enrichment score (NES) and FDR are shown in red. **(C)** Boxplots, comparing the mean z-scores for respiratory electron transport chain (ETC) genes across TCGA-SKCM tumors based on their NF1 mutation status. **(D)** Stage of tumor diagnosis in NF1$^{LoF}$ or NF1$^{LoF}$ and NF1$^{Mutant}$ patients based on their ETC expression levels. Tumors for each NF1 cohort were stratified into two subgroups, ETC$^{High}$ and ETC$^{Low}$, based on the top and bottom quartiles. Statistical significance ETC expression in NF1$^{LoF}$ patients versus NF1$^{Mutant}$ or NF1$^{Other}$ was determined by two-sided, two-sample t-test. Statistical significance was determined with Chi-squared test. **(E)** Dose-dependent changes in normalized growth rates (a.k.a. DIP rates) calculated by dividing the 7-day average net growth rate for inhibitor-treated cells to that measured for vehicle (DMSO)-treated cells across $NF1^{LoF}$ cell lines. The average net growth rates for each condition were calculated from measurements of live cell count (across two replicates) at four timepoints (including 1, 3, 5, and 7 days). For combination, the concentration of one inhibitor was varied between 0 and the indicated $C_{max}$ value in the presence of a fixed concentration of a second inhibitor. **(F)** Area over curve (AOC) of MTX-216, R406, or IACS-010759 alone (top) or in combination with trametinib (bottom) across $NF1^{LoF}$ melanoma cell lines. AOC value of 0 indicates no drug response. Statistical significance was determined by two-sided, paired-sample *t* test.

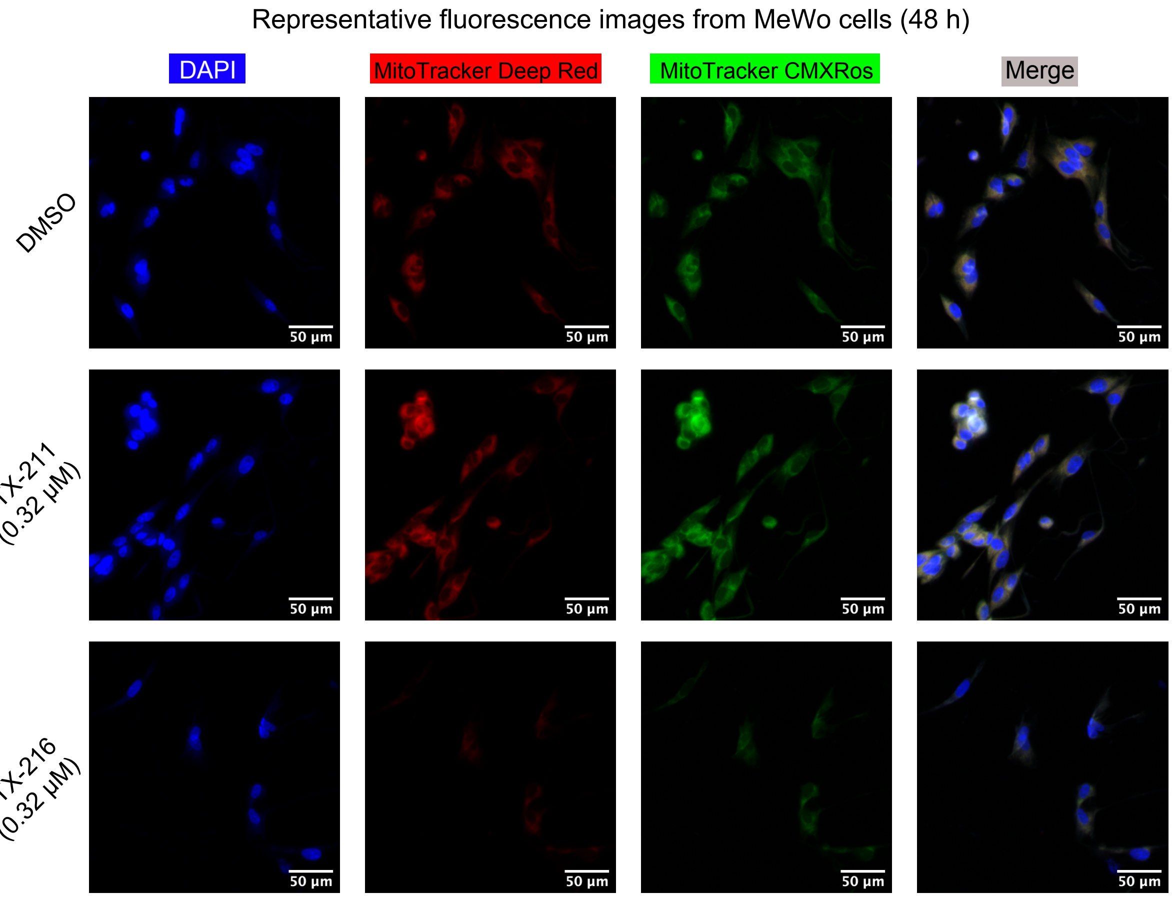


**Supplementary Figure S7. MTX-216 more potently suppresses mitochondrial mass and function compared to MTX-211.** Representative multiplexed fluorescent images of MitoTracker Deep Red and MitoTracker CMXRos, co-stained in MeWo cells following 48 h treatment with MTX-211 (0.316 µM), MTX-216 (0.316 µM), or DMSO. Each experiment was repeated three times independently with similar results. Scale bars represent 50 µm.
